# A selective bottleneck during host entry drives the evolution of new legume symbionts

**DOI:** 10.1101/2022.03.03.482760

**Authors:** Ginaini Grazielli Doin de Moura, Saida Mouffok, Nil Gaudu, Anne-Claire Cazalé, Marine Milhes, Tabatha Bulach, Sophie Valière, David Roche, Jean-Baptiste Ferdy, Catherine Masson-Boivin, Delphine Capela, Philippe Remigi

## Abstract

During the emergence of new host-microbe symbioses, multiple selective pressures-acting at the different steps of the microbial life cycle–shape the phenotypic traits that jointly determine microbial fitness. However, the relative contribution of these different selective pressures to the adaptive trajectories of microbial symbionts are still poorly known. Here we characterized the dynamics of phenotypic adaptation and its underlying genetic bases during the experimental evolution of a plant pathogenic bacterium into a legume symbiont. We observed that fast adaptation was predominantly driven by selection acting on competitiveness for host entry, which outweighed selection acting on within-host proliferation. Whole-population sequencing of evolved bacteria revealed that phenotypic adaptation was supported by the continuous accumulation of new mutations and the sequential sweeps of cohorts of mutations with similar temporal trajectories. The identification of adaptive mutations within the fixed mutational cohorts showed that all of them improved competitiveness for host entry, while only a subset of those also improved within host proliferation. Computer simulations predict that this effect emerges from the presence of a strong selective bottleneck at host entry. Together, these results show how selective bottlenecks can alter the relative influence of selective pressures acting during bacterial adaptation to multistep infection processes.

## Main

Many bacterial lineages have evolved the capacity to establish symbiotic associations, either beneficial, neutral or parasitic, with eukaryotic hosts. These interactions form a dynamic continuum along which bacteria can move^1^. Lifestyle changes may arise due to ecological (resource availability, host environment changes or host shifts) or genomic (mutations, acquisition of new genetic material) modifications. These new interactions are often initially sub-optimal for the bacterial partner^2–4^, which may then adapt to the selective pressures associated with its new life-cycle. For instance, horizontally-transmitted microbes alternate between at least two different habitats, the host and the environment, where they will face a variety of constraints and selective pressures^5,6^. Biphasic life cycles therefore require bacteria not only to be able to enter and exit their hosts, but also to replicate and persist within each habitat, which entails adapting to abiotic stressors, host immunity, competitors, as well as being able to use specific nutrient sources. However, the relative influence of the different selective pressures on the dynamics and the trajectory of bacterial adaptation to a new interaction is poorly known.

Rhizobia are examples of facultative host-associated bacteria that can either live freely in soil or in symbiotic mutualistic associations with legume plants^7^. In most legumes, rhizobia penetrate the root tissue through the formation of so-called infection threads (ITs). Most of the time, only one bacterium attaches to the root hair and initiates the formation of ITs^8^, thus creating a strong population bottleneck between the rhizosphere and internal root tissues. Bacteria then divide clonally within ITs, while at the same time a nodule starts to develop at the basis of the infected root hair. Hundreds of millions of bacteria are then released from ITs inside the cells of the developing nodule and differentiate into nitrogen fixing bacteroids^9,10^. After several months, in nature, nodule senescence leads to the release of a part of bacterial nodule population in the surrounding soil. To fulfil their life cycle, rhizobia rely on the activity of numerous bacterial genes, allowing signal exchanges with the host plant, proliferation and metabolic exchanges within nodule cells, while maintaining free-living proficiency^7^. Somewhat surprisingly given their complex and specialized lifestyle, rhizobia evolved several times independently. Horizontal transfer of key symbiotic genes is a necessary, though often not sufficient, condition for a new rhizobium to emerge^11–14^. Additional steps of adaptation of the recipient bacterium, occurring during evolution under plant selection, are believed to be needed to actualize the symbiotic potential of emerging rhizobia^15^.

In a previous study, we used experimental evolution to convert the plant pathogen *Ralstonia solanacearum* into an intracellular legume symbiont^16^. Here, we analysed the dynamics of molecular evolution in 5 lineages evolved for 35 cycles of nodulation through whole-population genome sequencing and examined the selective and genetic bases of adaptation. Adaptation proceeded rapidly during the first cycles of evolution, and was underpinned by the fixation of successive cohorts of mutations within populations. We then identified adaptive mutations in two lineages and evaluated their effect on each symbiotic stage of the rhizobial life cycle. Our experimental data indicated that selection for nodulation competitiveness outweighs selection for multiplication within host. Computer simulations further showed that the selective bottleneck at host entry and the chronology of symbiotic events are critical drivers of this evolutionary pattern.

## Results

### Fast adaptation of new legume symbionts during the first cycles of evolution

To investigate the genetic and evolutionary conditions that promote the evolution of new rhizobia, we previously transferred a symbiotic plasmid (pRalta) from the rhizobium *Cupriavidus taiwanensis* LMG19424 into the plant pathogen *Ralstonia solanacearum* GMI1000. We obtained three nodulating ancestors, two infecting nodules intracellularly (CBM212, CBM349) and one infecting nodules extracellularly (CBM356)^14^, that we evolved in 18 parallel bacterial lineages for 16 serial cycles of nodulation on *Mimosa pudica* plants^16^. Here we continued the evolution of five of these lineages (referred to as lineages B, F, G, K, and M) until cycle 35 (Figure 1a, Figure 1 – figure supplement 1 and Supplementary file 1). After 35 cycles, no nitrogen fixation sustaining plant growth was observed (Figure 1 – figure supplement 2). Therefore, in this manuscript, we focussed exclusively on the analysis of changes in bacterial *in planta* fitness (estimated for each strain as the frequency of bacteria present in nodules relative to a competing reference strain). To analyse the dynamics of fitness changes over time, we compared the relative fitness of the nodulating ancestors and evolved clones from cycles 16 and 35 by replaying one nodulation cycle in competition with the natural symbiont *C. taiwanensis*. Fitness trajectories were similar in the five lineages. While the three nodulating ancestors were on average 10^6^ times less fit than *C. taiwanensis*, fitness improved very quickly during the first 16 cycles, then slower until cycle 35 (Figure 1b). On average, evolved clones isolated at cycle 16 and cycle 35 were 47 and 19 times less fit than *C. taiwanensis*, respectively (Figure 1b).

**Figure 1:**
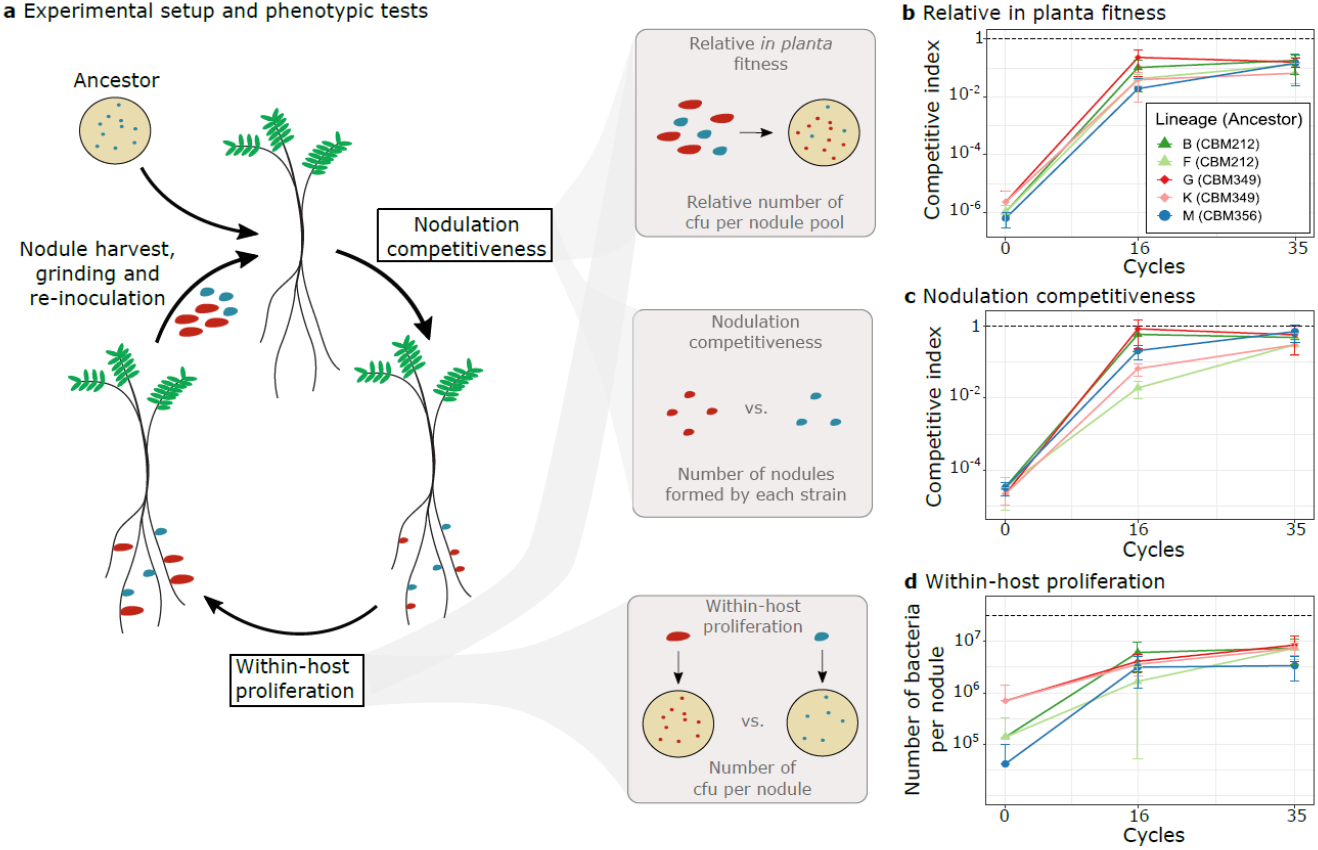
Evolution of the symbiotic properties of *Ralstonia* clones along evolution cycles. a, Left: overview of the evolution cycles showing the symbiotic steps determining bacterial fitness (nodulation competitiveness and within-host proliferation) and human intervention (nodule harvest, grinding and re-inoculation). Nodule colors reflect bacterial genotypes: blue represents the ancestor, and red represents a mutant with increased fitness. Right: schematic representation of the main phenotypic measurements performed in this work, where the symbiotic phenotypes of one evolved strain can be compared to those of a reference strain (e.g. the reference symbiont *C. taiwanensis)*. Competitive indexes for *in planta* fitness are calculated as the ratio of evolved vs. reference clones in nodule bacterial populations normalized by the ratio of strains in the inoculum. Competitive indexes for nodulation competitiveness are calculated as the ratio of nodules formed by each strain normalized by the inoculum ratio. Within-host proliferation is measured in independent single inoculations of each strain. b-d, Relative *in planta* fitness (b), nodulation competitiveness (c) and within-host proliferation (d) of nodulating ancestors (cycle 0) and evolved clones isolated from cycles 16 and 35 were compared to *C. taiwanensis*. Values correspond to means ± standard deviations. Data were obtained from at least three independent experiments. For each experiment, nodules were harvested from 10 plants (b), 20 plants (c) and 6 plants (d). The sample size (n) is equal to n=3 (b,c) or comprised between n=15–18 (d). Raw data are available in Figure 1 – source data 1. cfu: colony-forming units. **Figure 1 – figure supplement 1:** Experimental evolution of *Ralstonia solanacearum* GMI1000 pRalta through serial cycles of plant (*Mimosa pudica*) inoculation-isolation of nodule bacteria. **Figure 1 – figure supplement 2:** Dry weights of *M. pudica* plants inoculated with cycle 35 evolved clones. **Figure 1 – source data 1:** Raw data obtained for the phenotypic characterization of *Ralstonia* evolved clones compared to *C. taiwanensis*. **Figure 1 – figure supplement 2 – source data 1:** Raw data obtained for the measurements of the dry weights of *M. pudica* plants inoculated with cycle 35 clones.

Because bacterial fitness in our system depends on both the capacity of strains to enter the host and induce nodule formation (nodulation competitiveness) and to multiply within these nodules (within-host proliferation), we analysed how each of these two fitness components changed during the experiment (Figure 1c and d). In the five lineages, evolutionary trajectories of nodulation competitiveness resembled that of fitness with fast improvement during the first 16 cycles, and a slow-down during the last cycles. On average, nodulating ancestors were 5×10^4^ times less competitive than *C. taiwanensis*, while cycle 16 and cycle 35 evolved clones were only 17 and 4 times less competitive than *C. taiwanensis*, respectively (Figure 1c). Moreover, evolved clones B16, B35, G16, G35 and M35 were not statistically different from *C. taiwanensis* in terms of nodulation competitiveness. Within-host proliferation also improved mostly during the first 16 cycles in all lineages. The rate of improvement then slowed down in two lineages and continued to increase significantly in the three others. As previously published, the three nodulating ancestors display different capacities to infect nodules^14,16^, the extracellularly infective ancestor CBM356 being the less infective (750 times less than *C. taiwanensis)* and the intracellularly infective CBM349 being the most infective (45 times less than *C. taiwanensis)*. On average, cycle 16 and cycle 35 evolved clones were 8 and 5 times less infective than *C. taiwanensis* (Figure 1d).

Altogether, the symbiotic properties of evolved clones improved rapidly during the first cycles of the evolution experiment and then adaptation slowed down. Both symbiotic traits, nodulation competitiveness (*i.e*. host entry) and within-host proliferation, improved greatly. However, gains in nodulation competitiveness (average factor of 11,500 between ancestors and cycle 35 clones) were *ca*. 150 times higher than gains in proliferation (average factor of 74 between ancestors and cycle 35 clones). Moreover, although the difference in nodulation competitiveness between *C. taiwanensis* and the nodulating ancestors was much more important (>10^4^ fold) than the difference in proliferation (<10^3^ fold), nodulation competitiveness of evolved clones reached the level of *C. taiwanensis* in three lineages (B, G and M) while within host proliferation remained lower than *C. taiwanensis* in all of them after 35 cycles.

### The dynamics of molecular evolution is characterized by multiple selective sweeps of large mutational cohorts

The rapid adaptation in this experiment was unexpected given that strong population bottlenecks, such as the one occurring at the nodulation step, limit effective population size at each cycle (see Supplementary file 1) and are generally known to limit the rate of adaptive evolution^17^. To describe the dynamics of evolution at the molecular level, we performed whole-population sequencing of evolved lineages using the Illumina sequencing technology. Populations of the five lineages were sequenced every other cycle until cycle 35 with a minimum sequencing coverage of 100× and a median of 347× (Supplementary file 2). We detected a very large number of mutations in all sequenced populations (a total of 4114 mutations, between 382 and 1204 mutations per lineage with 23.6 new mutations per cycle on average) (Supplementary file 3 and Figure 2 – source data 1). This high number of mutations is a result of transient hypermutagenesis occurring in the rhizosphere^18^. Mutations accumulated in the populations throughout the experiment, showing no sign of slowing down until the end of the experiment, which suggests that no anti-mutator mutations established in these populations as we might have expected^19^. We evaluated genetic parallelism among mutations detected above a frequency of 5% in the populations by computing G scores^20^, a statistics used to point out genes that could be mutated more often than expected by chance (Supplementary file 4). In spite of high mutation rates that may obscure signals of genetic convergence, we detected signatures of parallelism at the gene level among our list of mutations (observed sum of G scores of 7,540.5, compared to a mean sum of 5,401.01 after 1,000 randomized simulations, Z = 31.23, *P* < 10^−200^). The randomized simulations identified 171 genes with a higher number of mutations than expected by chance (Bonferroni adjusted *P* value<0.01, Supplementary file 4).

To simplify the analysis of mutational trajectories, we then focused on the 819 mutations (out of a total of 4414 detected mutations) that rose above a frequency of 30%. These mutations did not show independent trajectories. Instead, we observed that groups of mutations arose synchronously and showed correlated temporal trajectories, enabling us to cluster them in cohorts^21,22^ (Figure 2 and Supplementary file 5). Cohorts increased in frequency with variable speed, some being fixed within 1 to 3 cycles (representing *ca*. 25–75 bacterial generations) while others reached fixation in up to 21 cycles (representing more than 500 bacterial generations). Fixed cohorts can be very large (up to 30 mutations) (Supplementary file 5), which suggests hitchhiking of neutral or slightly deleterious mutations with one or several beneficial mutations acting as driver (or co-drivers) of the cohort^21,23^. Among the mutations that rose above 30% frequency at some stage of the experiment, 35% later declined until extinction. This is indicative of clonal interference, *i.e*. the co-occurrence of multiple subpopulations competing against each other within populations^24^, which can readily be observed on Muller plots for lineages B and G (Figure 2 – figure supplement 1).

Overall, in these 5 lineages, the pattern of molecular evolution was characterized by a steady accumulation of mutations along the successive cycles and the formation of mutational cohorts, some of which containing a large number of mutations. Despite strong population bottlenecks at the root entry, mutation supply was not limiting as evidenced by the co-occurrence of competing adaptive mutations. Yet, multiple (and sometimes rapid) selective sweeps occurred throughout the 35 cycles in all lineages, suggesting that strong selection is acting on these populations.

### Large fixed cohorts contain multiple adaptive mutations

In our system, strong nodulation bottlenecks lead to a small effective population size and to the possibility that genetic drift might be responsible for the observed allelic sweeps. To determine whether drift or adaptation are causing these sweeps, we searched for adaptive mutations in fixed (or nearly fixed, >90% frequency) cohorts from the two lineages that have the highest symbiotic fitness after 35 cycles: B and G. To do so, we introduced (by genetic engineering) individual mutations from each cohort of interest into an evolved clone carrying all (or, when not available, almost all) previously fixed cohorts. This allowed us to test the fitness effect associated with each mutation in a relevant genetic background, taking into account possible epistatic effects arising from mutations that were previously acquired in this clone. Competitive index (ratio of strains carrying the mutant vs. the wild-type alleles in nodule populations normalized by the inoculum ratio) were measured for 44 mutations belonging to 14 cohorts from the two lineages (Figure 3a and b). Twenty-eight mutations significantly increased the fitness of evolved clones and were thus beneficial for symbiosis while 11 mutations were neutral and 5 were slightly deleterious for symbiosis (Figure 3a and b, Table 1 and Supplementary file 6). Among the 44 genes targeted by the reconstructed mutations, 8 have a higher number of mutations than expected by chance (Supplementary file 4) and adaptive mutations were identified in 6 of them. Several interesting results emerge from this data set. First, some mutations show very strong adaptive effects, improving more than 100 times the fitness of bacteria, in particular during the first cycles of evolution. Second, many cohorts carry more than 1 adaptive mutation: six cohorts contain between 2 and 7 adaptive mutations. Third, 35% of adaptive mutations were found to be synonymous mutations, confirming that these mutations can play a significant role in adaptive evolution^25^. All adaptive synonymous mutations except one converted frequently used codons in *R. solanacearum* into unusual ones (Supplementary file 7), which may explain their functional effects. Moreover, when inspecting gene annotations, we found that the 28 beneficial mutations target various biological functions (Table 1). Strikingly, nearly 40% affect regulatory functions such as global transcription regulators (*efpR* and Rsc0965)^26^, putative quorum quenching (RSp0595), signal transduction (RSc1598), unknown transcription regulator (RSc0243), protein dephosphorylation (Rsp1469), protein folding (*ppiB*), degradosome (*rhlE1*), and DNA methylation (RSc1982). Another significant proportion of mutations (32%) affect metabolism and transport genes (*gloA*, *phbB*, RSc0194, RSc2277, Rsp1116, RSp1417, RSp1422, RSp1590 and pRALTA_0398) and 3 mutations are located in genes belonging to the type VI secretion system (T6SS) operon (*tssJ*, RSp0759, Rsp0769). The observed genetic architecture (*i.e*. the large number of genes and molecular functions targeted by adaptive mutations) indicates that symbiotic traits are complex and constitute a large mutational target for adaptive evolution.

**Figure 2:**
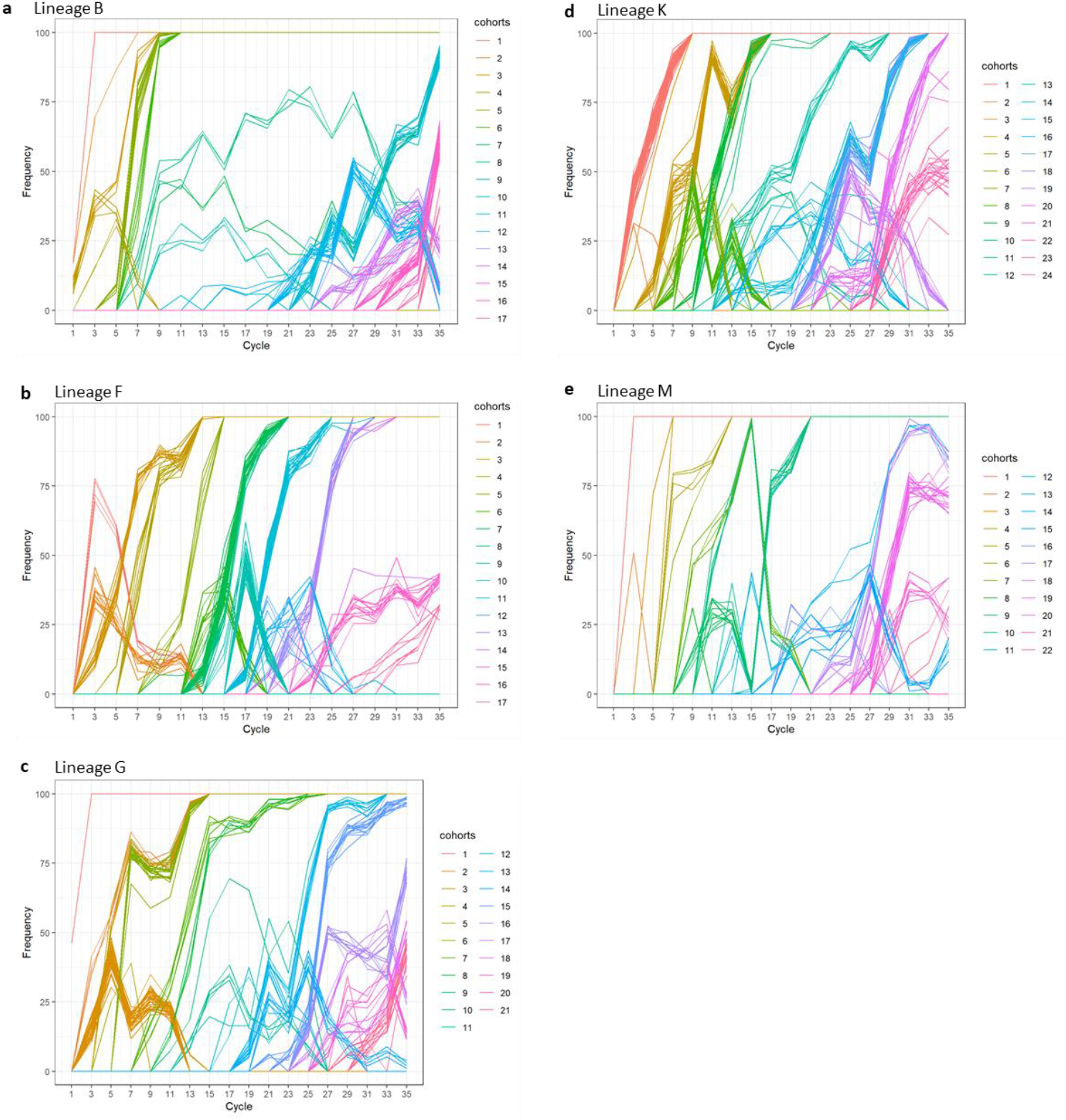
Dynamics of molecular evolution. Allele frequency trajectories of mutations that attained a frequency of 30% in at least one population of the B, F, G, K and M lineages (**a**,**b**,**c**,**d**,**e**). Mutations with similar trajectories were clustered in cohorts and represented by different colors. For simplicity, mutations travelling alone were also called cohorts. **Figure 2 – figure supplement 1:** Evolution of population composition over time in lines B and G. **Figure 2 – source data 1:** Mutations identified in the evolved populations from all lineages **Figure 2 – source data 2:** Mutations present in the evolved clones

**Figure 3:**
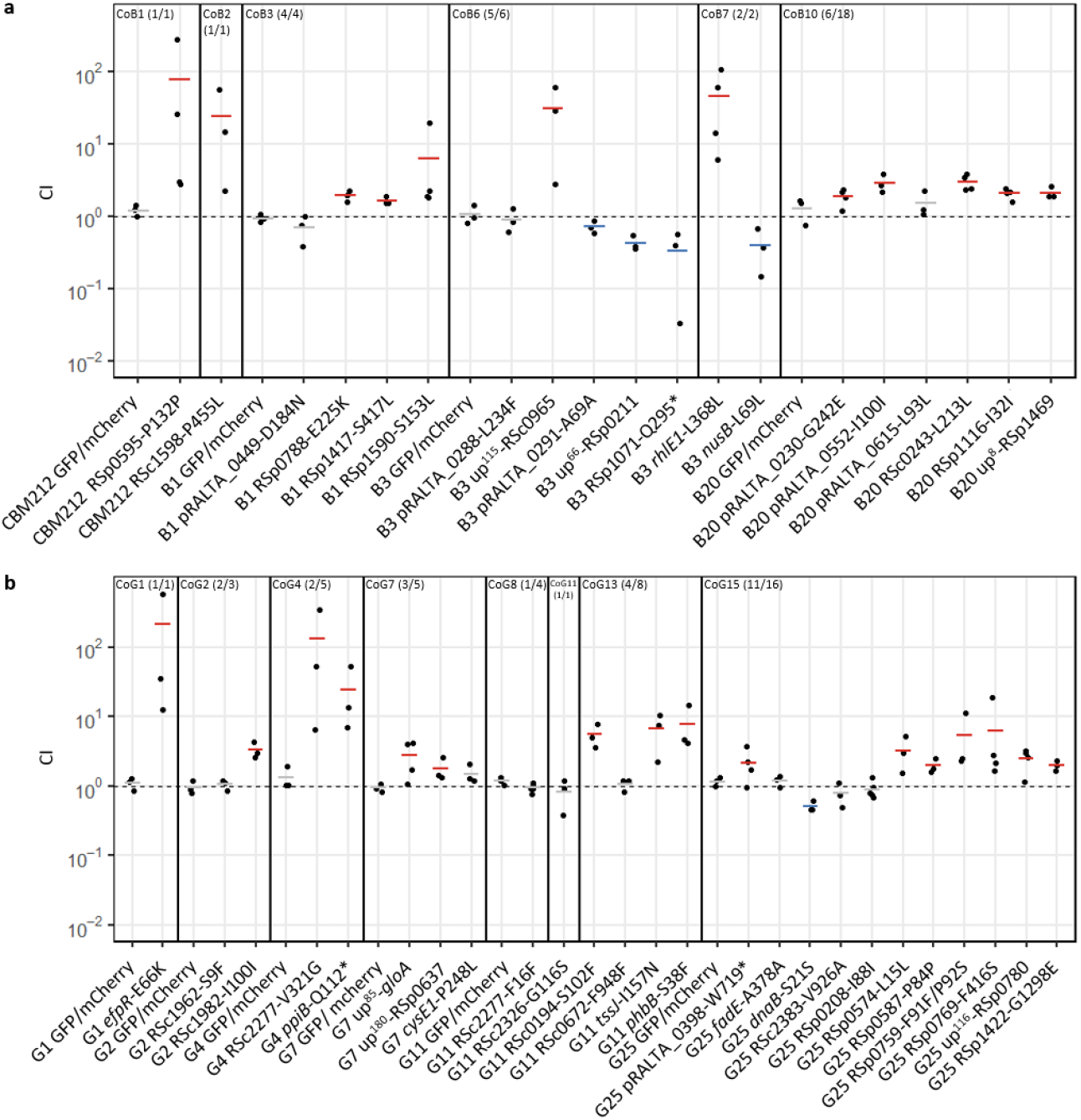
*In planta* fitness of reconstructed mutants from lineages B and G. *In planta* relative fitness of evolved clones carrying reconstructed mutations from fixed mutational cohorts identified in lineages B (a) and G (b). Evolved clones in which mutations were reconstructed are indicated at the bottom of the graph. Vertical lines separate the different mutational cohorts. Cohort number and the number of tested mutations from the cohort on the total number of mutations present in the cohort are indicated in brackets. Competitive indexes (CI) were calculated as the ratio of the mutant strain on the isogenic parental strain in bacterial nodule populations normalized by the inoculum ratio. GFP/mCherry correspond to control co-inoculation experiments of strains derived from the same evolved clone labelled with different fluorophore either GFP or mCherry. Horizontal segments correspond to mean values of CI. Red segments indicate significantly beneficial mutations, gray segments indicate neutral mutations and blue segments indicate significantly deleterious mutations (*P*<0.05, *t*-test). Data were obtained from 3 to 5 independent experiments. The sample size (n) is comprised between n=3–5. Each CI value was obtained from pools of 40 to 227 nodules harvested from 10 plants. **Figure 3 – source data 1**: Raw data obtained for the characterization of the *in planta* fitness of evolved clones carrying fixed mutations.

**Table 1:** Symbiotic phenotypes of beneficial mutations improving *in planta* fitness.

| Lineage | Gene ID<sup>a</sup> | Gene name | Mutation | Type of mutation<sup>b</sup> | Description | Cohort | Mean CI for *in planta* fitness<sup>c</sup> | Mean CI for plant culture medium colonization<sup>d</sup> | Mean CI for rhizosphere colonization<sup>d</sup> | Mean CI for nodulation competitiveness<sup>c</sup> | Mean gain factor in within-host proliferation<sup>e</sup> | Symbiotic phenotypes<sup>f</sup> | G score<sup>g</sup> | Adjusted P value<sup>h</sup> |
| --- | --- | --- | --- | --- | --- | --- | --- | --- | --- | --- | --- | --- | --- | --- |
| B | RSp0595 |  | P132P | Syn | Putative N-acyl-homoserine lactonase | CoB1 | **77** | 1 | 1 | **5.2** | 1.6 | Nod+ | 2.80 | 1 |
|  | RSc1598 |  | P455L | Nonsyn | Transmembrane sensor histidine kinase | CoB2 | **24.2** | 1 | 1.4 | **4.7** | **6.4** | Nod+ Pro+ | 1.04 | 1 |
|  | RSp0788 |  | E225K | Nonsyn | Carbamoyl-transferase | CoB3 | **1.9** | 1 | 1.1 | **2** | 1.2 | Nod+ | -0.09 | 1 |
|  | RSp1417 |  | S417L | Nonsyn | Multidrug efflux transmembrane protein | CoB3 | **1.6** | 0.8 | 1.4 | **3.9** | 0.9 | Nod+ | 12.47 | **2.58E-04** |
|  | RSp1590 |  | S153L | Nonsyn | FAD-dependent oxireductase | CoB3 | **6.3** | 1 | 0.9 | **2.4** | 0.6 | Nod+ | 4.10 | 1 |
|  | up<sup>115</sup>-RSc0965 |  | G/A | Intergenic | Unknown function, negatively regulates *efpR* expression | CoB6 | **31.1** | 0.9 | **2.9** | **8.2** | **2.6** | Rhizo+ Nod+ Pro+ | nd | nd |
|  | RSc0539 | *rhIE1* | L368L | Syn | ATP-dependent RNA helicase protein | CoB7 | **46.2** | 1.1 | **4** | **28.2** | **2.4** | Rhizo+ Nod+ Pro+ | 2.88 | 1 |
|  | RSc0243 |  | L213L | Syn | LysR-type transcription regulator protein | CoB10 | **3** | 1.3 | 1.4 | **3.9** | 1.1 | Rhizo+ Nod+ | 1.16 | 1 |
|  | RSp1116 |  | I32I | Syn | Putative low specificity L-threonine aldolase | CoB10 | **2.1** | 1.2 | 1.2 | **2.1** | 0.9 | Nod+ | 0.84 | 1 |
|  |  |  | L33F | Nonsyn |  |  |  |  |  |  |  |  |  |  |
|  | up<sup>8</sup>-RSp1469 |  | C/T | Intergenic | Putative tyrosine specific protein phosphatase | CoB10 | **2.1** | 0.9 | 1 | **2** | 1.0 | Nod+ | nd | nd |
|  | pRALTA_0230 |  | G242E | Nonsyn | Putative ATP-dependent DNA helicase | CoB10 | **1.5** | 1.5 | 1.5 | **2.1** | 0.9 | Nod+ | 6.34 | 1 |
|  | pRALTA_0552 |  | I100I | Syn | Pseudogene, putative membrane bound hydrogenase | CoB10 | **2.9** | 1.2 | 1.4 | **2.3** | 0.8 | Nod+ | nd | nd |
| G | RSc1097 | *efpR* | E66K | Nonsyn | Transcriptional regulator | CoG1 | **212.1** | 1.3 | **4** | **35** | **3.7** | Rhizo+ Nod+ Pro+ | 3.17 | 1 |
|  | RSc1982 |  | I100I | Syn | DNA methyltransferase | CoG2 | **3.3** | 3.1 | 1.3 | **4.0** | 0.7 | Nod+ | 1.44 | 1 |
|  | RSc1164 | *ppiB* | Q112* | Nonsense | Peptyl-prolyl-cis/trans isomerase | CoG4 | **24.5** | 1 | 1.0 | **5.0** | 1.4 | Nod+ | 2.26 | 1 |
|  | RSc2277 |  | V321G | Nonsyn | Putative transporter protein | CoG4 | **133.9** | 9.3 | 10.1 | **58** | **2.5** | Nod+ Pro+ | 36.95 | **4.68E-50** |
|  | up<sup>85</sup>-RSc0520 | *gloA* | T/C | Intergenic | Lactoylglutathione lyase | CoG7 | **2.8** | 1.2 | **1.5** | **2.1** | 1.1 | Nod+ | nd | nd |
|  | up<sup>180</sup>-RSp0637 |  | G/C | Intergenic | Hypothetical protein | CoG7 | **1.8** | 1 | 1.2 | **2.6** | 0.9 | Nod+ | nd | nd |
|  | RSc0194 |  | S102F | Nonsyn | Zinc-dependent alcohol dehydrogenase | CoG13 | **5.5** | 1.1 | 1.6 | **3.1** | 1.4 | Nod+ | 4.67 | 1 |
|  | RSc1633 | *phbB* | S38F | Nonsyn | Acetoacetyl-CoA reductase | CoG13 | **7.9** | 2 | 2.1 | **3.3** | 1.7 | Rhizo+ Nod+ Pro+ | 1.62 | 1 |
|  | RSp0741 | *tssJ* | I157N | Nonsyn | Type VI secretion system, lipoprotein TssJ | CoG13 | **6.7** | 1 | 1.1 | **2.9** | 1.7 | Nod+ | 6.99 | 1 |
|  | RSp0574 |  | L15L | Syn | Hypothetical protein | CoG15 | **3.2** | 1.1 | 1.3 | **2.1** | **1.6** | Nod+ Pro+ | 4.36 | 1 |
|  | RSp0587 |  | P84P | Syn | Putative signal peptide hypothetical protein | CoG15 | **2** | 1.1 | 0.8 | **2.6** | 1.3 | Nod+ | 28.74 | **1.04E-46** |
|  | RSp0759 |  | F91F | Syn | Putative type VI secretion-associated protein | CoG15 | **5.4** | 0.7 | 0.9 | **2.5** | 1.4 | Nod+ | 1.00 | 1 |
|  |  |  | P92S | Nonsyn |  |  |  |  |  |  |  |  |  |  |
|  | RSp0769 |  | F416S | Nonsyn | Putative type VI secretion-associated protein | CoG15 | **6.3** | 1.1 | 1.3 | **1.6** | **1.8** | Nod+ Pro+ | 17.20 | **3.98E-10** |
|  | up<sup>116</sup>-RSp0780 |  | C/T | Intergenic | Hypothetical protein | CoG15 | **2.5** | 1.1 | 1.1 | **3** | 1.1 | Nod+ | nd | nd |
|  | RSp1422 |  | G1298E | Nonsyn | Non-ribosomal peptide synthase | CoG15 | **2** | 1.2 | 1.2 | **2.4** | 1.6 | Nod+ | 18.28 | **2.79E-03** |
|  | pRALTA_0398 |  | W719* | Nonsense | Phosphoenol pyruvate synthase | CoG15 | **2.2** | 0.8 | 0.8 | **1.6** | **1.6** | Nod+ Pro+ | 151.77 | **0** |

<sup>a</sup>up<sup>x</sup>, intergenic mutations located x nucleotides upstream the gene indicated. <sup>b</sup>Syn, synonymous mutations. Nonsyn, non synonymous mutations. <sup>c</sup>Mean values of competitive indexes (CI) obtained from 3 to 4 independent experiments. Figures in bold are statistically different from 1 (Student *t*-test *P* < 0.05). <sup>d</sup>Mean values obtained from at least 15 measurements from 3 to 4 independent experiments. Figures in bold are statistically significant (Wilcoxon test, *P* < 0.05). <sup>e</sup>Mean values obtained from at least 15 measurements from 3 to 4 independent experiments. Figures in bold are statistically different from 1 (Wilcoxon test, *P* < 0.05 with the Benjamini-Hochberg correction). <sup>f</sup>Rhizo+, improvement in rhizosphere colonization. Nod+, improvement in nodulation competitiveness. Pro+, improvement in within-host proliferation. <sup>g</sup>G scores were calculated as proposed by Tenaillon et al. (2016). nd, G scores were not determined for mutations in pseudogenes or intergenic regions. <sup>h</sup>P-values associated with G scores were adjusted using a Bonferroni correction to evaluate genetic parallelism.

### Adaptive mutations predominantly improve nodulation competitiveness

Next, we measured the effect of each adaptive mutation on nodulation competitiveness (reflecting host entry) and within-host proliferation in order to uncover which of these two symbiotic traits can explain the observed fitness gain. All mutations increasing fitness improved nodulation competitiveness of evolved clones (Figure 4a and b). Gains in nodulation competitiveness ranged from 1.6 to 58-fold, with three mutations providing the highest gains: RSc2277-V321G (58-fold), *efpR-E66K* (35-fold) and *rhlEl-L368L* (28-fold). By contrast, within-host proliferation was improved by only a subset of adaptive mutations (Figure 5a and b). Only 3 mutations from lineage B (RSc1598-P455L, up^115^-RSc0965 and *rhlEl-L368L*) and 7 mutations from lineage G (*efpR*-E66K, RSc2277-V321G, *phbB-S38F*, pRALTA_0398-W719^*^, RSp0574–L15L, RSp0769-F416S and RSp1422-G1298E) significantly improved this fitness component. Moreover, gains in *in planta* proliferation were generally lower than gains in nodulation competitiveness and ranged from 1.6 to 6.4-fold. The highest proliferation gains were produced by the five mutations RSc1598-P455L (6.4-fold), *efpR*-E66K (3.7-fold), up^115^-RSc0965 (2.6-fold), RSc2277-V321G (2.5 fold) and *rhlEl*-L368L (2.4-fold), three of which also produced the highest nodulation gains.

**Figure 4:**
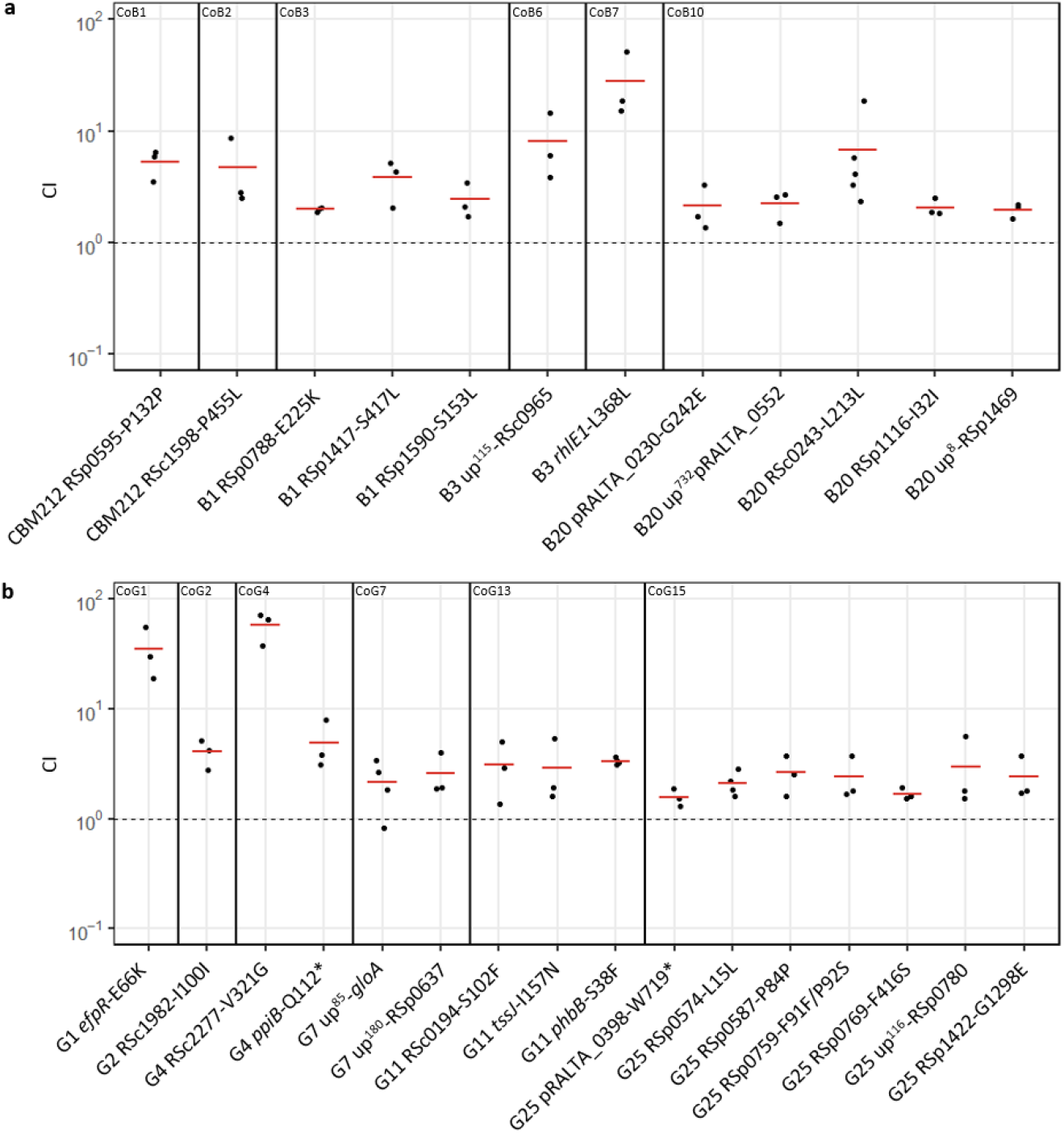
Nodulation competitiveness of adaptive mutants. Nodulation competitiveness effect of adaptive mutants from lineages B (**a**) and G (**b**). Evolved clones in which mutations were reconstructed are indicated at the bottom of the graph. Vertical lines separate the different mutational cohorts. Cohort number and the number of tested mutations from the cohort on the total number of mutations present in the cohort are indicated in brackets. CI were calculated as the ratio of the number of nodules formed by the mutant strain on the number of nodules formed by the isogenic parental strain normalized by the inoculum ratio. Horizontal segments correspond to mean values of CI. Red segments indicate significantly beneficial mutations (*P*<0.05, *t*-test). Data were obtained from 3 to 4 independent experiments. The sample size (n) is comprised between n=3–4. Each CI value was obtained from 95–98 nodules harvested from 20 plants. **Figure 4 – figure supplement 1:** Survival of reconstructed mutants from lineages B and G in the Jensen medium and rhizosphere. **Figure 4 – source data 1:** Raw data obtained for the characterization of nodulation competitiveness of evolved clones carrying fixed mutations. **Figure 4 – figure supplement 1 – source data 1**: Raw data obtained for the Jensen culture medium colonization and rhizosphere colonization assays.

**Figure 5:**
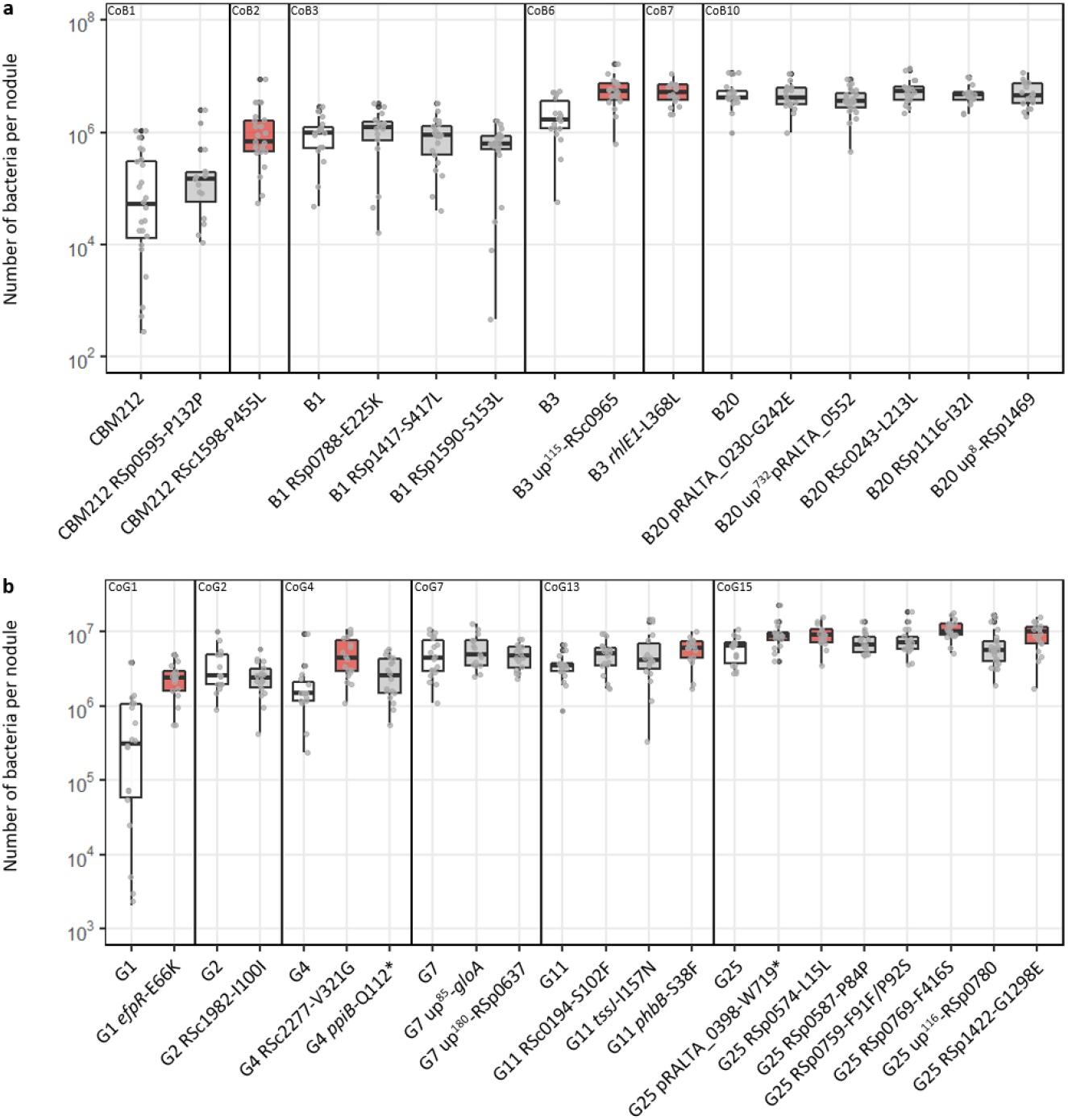
Within-host proliferation of adaptive mutants and corresponding isogenic parental clones. Distribution of the number of bacteria per nodule recovered for adaptive mutants from lineage B (a) and G (b) and their corresponding isogenic parental clones at 21 dpi. Rectangles span the first quartiles to the third quartiles, bold segments inside the rectangle show the median, unfiled circles represent outliers, whiskers above and below each box show the minimum and the maximum in the absence of suspected outlying data. Red boxes indicate the mutations that significantly improved infectivity compared to the parental evolved clone (P<0.05, Wilcoxon test). Data were obtained from 3 to 5 independent experiments. For each experiment, nodules were harvested from 6 plants. The sample size (n) is comprised between n=15–24. **Figure 5 – source data 1:** Raw data obtained for the characterization of the capacity of evolved clones carrying fixed mutations to proliferate within the host.

To investigate whether improvements in nodulation competitiveness could be due to a better colonization of the plant culture medium or rhizosphere, we measured the survival of the 28 mutants in competition with their isogenic parental strain in these two compartments. Results from these assays showed that none of the adaptive mutations improved bacterial ability to colonize the culture medium and only 4 of them, including up^115^-RSc0965, *efpR*-E66K and *rhlEl*-L368L, improved the colonization of the rhizosphere (Figure 4 – figure supplement 1) although by smaller factors than improvements in nodulation competitiveness. These results showed that improvements in nodulation competitiveness were generally not associated with a better colonization of the culture medium or of the rhizosphere, indicating that host entry is mainly controlled directly by the plant and is the dominant selective force driving the adaptation of legume symbionts in this evolution experiment.

### Evolutionary modelling predicts that selection for host entry drives adaptation

The preferential selection of nodulation competitiveness in our experiment prompted us to investigate the origin of this phenomenon. In particular, we wondered if stronger selection for host entry over within-host proliferation could be a general feature of symbiotic life cycles, or, instead, if it is more likely to be a specific property of our experimental system that may arise due to genetic constraints on symbiotic traits and/or specific features of the selective regime. We used computer simulations to model the evolution of bacterial populations that cycle between two compartments: the external environment (rhizosphere) and the host tissues (root nodules). In our model, bacteria accumulate mutations in the rhizosphere due to transient hypermutagenesis^18^ while we consider that the bacterial population remains clonal within nodules^18^ (see Supplementary file 8 for full details on the model and Figure 6 – figure supplement 1). These assumptions are relevant not only to rhizobium-legume interactions but also to other horizontally transmitted symbioses where the hosts are exposed to highly diverse environmental bacterial populations and accommodate more homogenous populations within their tissues^27–29^. In our experimental system, bacterial fitness is mediated by two phenotypic components: competitiveness to enter the host (resulting in nodulation) and within-host proliferation. Host entry imposes a very stringent selective bottleneck since (i) in most cases, each nodule is founded by one single bacterium^9,30^, (ii) there were usually between 100 and 300 nodules per lineage collected at each cycle (Supplementary file 1) while *ca*. 10^6^–10^7^ bacteria were inoculated at each cycle and (iii) bacterial genotypes differ in their competitive ability to form nodules (Figure 4a and b). Once inside the host, bacteria multiply to reach a carrying capacity that is directly proportional to their proliferation fitness, before returning to the external environment.

When following the evolution of bacterial phenotypes in populations founded by an ancestor with low initial fitness (10^−4^ fold compared to the theoretical optimum, for each phenotypic component), we observed, in most cases, a faster increase in nodulation competitiveness relative to within-host proliferation (Figure 6a). Accordingly, the distribution of mutations that were selected at the early steps of the adaptive process was biased towards stronger improvement of nodulation competitiveness over proliferation (Figure 6b, Low fitness domain). This dominance decreased, and was sometimes reverted, as populations progressed towards high fitness values. Decreasing the strength of the host entry bottleneck also reduced the dominance of early selection for nodulation competitiveness, until abolishing it for the largest bottleneck size tested (3000 nodules).

**Figure 6:**
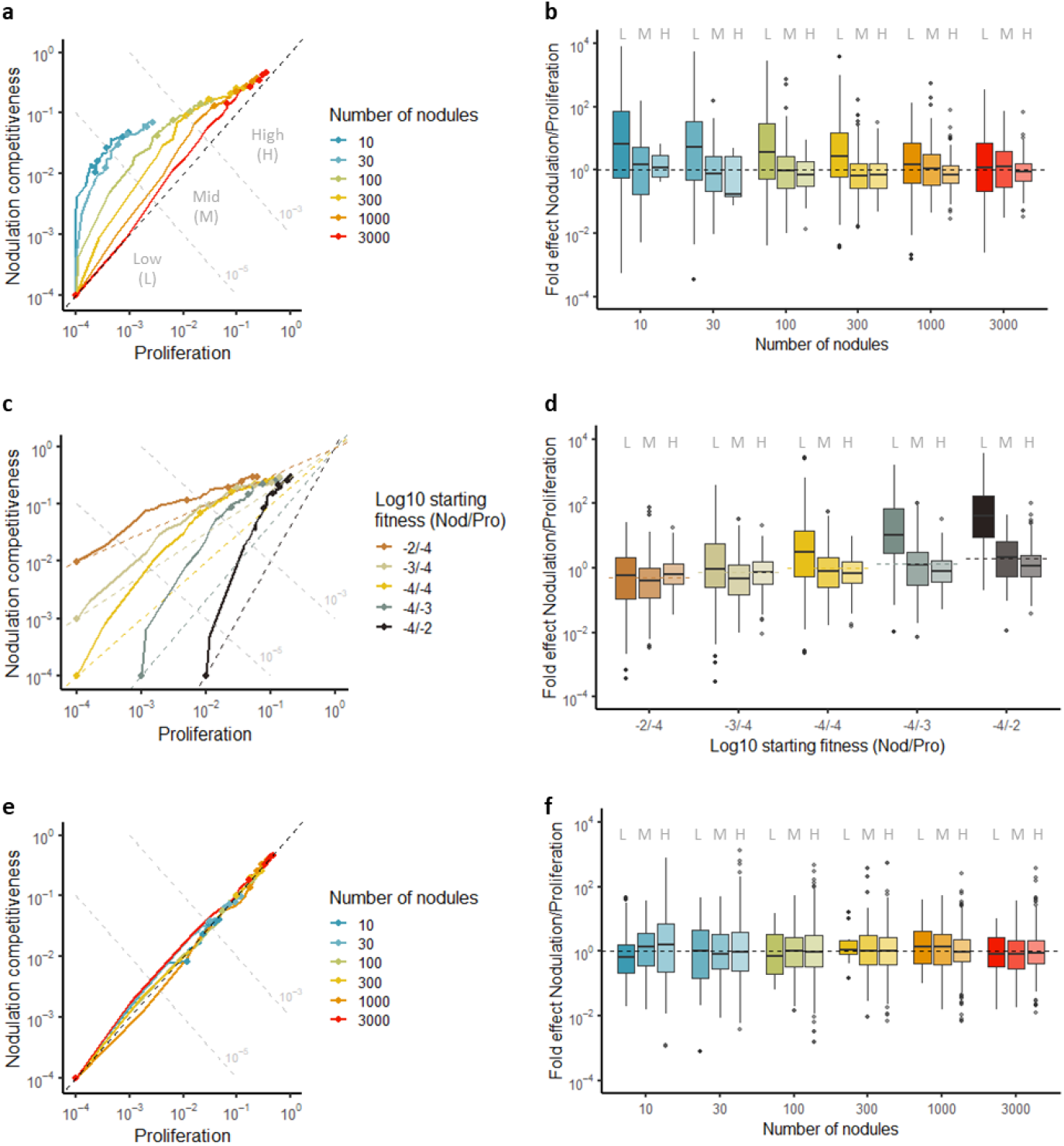
Relative strength of selection for nodulation competitiveness and within-host proliferation. **a,c,e**, Median fitness trajectories of 100 simulated populations evolving under different sizes of host entry bottleneck (10 to 3000 nodules). Points indicate fitness values at cycles 0, 10, 20, 30, 40 and 50. Dotted grey lines represent iso-fitness lines (defined as the product of nodulation competitiveness and within-host proliferation values) used to delineate 3 fitness domains: Low (L; fitness < 10^−5^), Mid (M; 10^−5^ < fitness < 10^−3^), and High (H; fitness > 10^−3^). The black dotted line represents the diagonal, along which population would improve both phenotypic traits equally well. **b,d,f**, Fold effect (nodulation competitiveness effect divided by proliferation effect) of mutations that reached a frequency of at least 30% in simulated populations. Color shading indicates the fitness domain (Low, Mid or High) in which each group of mutations arose. Note that fold effects of mutations are indicated even when median fitness trajectories do not reach the High fitness domain (*e.g*. 10 and 30 nodule lines in **a**) because some individual replicate simulations (shown in Supplementary Figures) do reach this fitness domain. The effect of different parameters on adaptive trajectories was tested: the size of host entry bottleneck (**a, b**), the initial fitness values of the ancestor (**c, d**), and the size of population bottleneck under a scenario where the bottleneck occurs after bacterial clonal proliferation (**e, f**). **Figure 6 – figure supplement 1:** Schematic representation of the modelling framework. **Figure 6 – figure supplement 2:** Distribution of fitness effects of new mutations for each of the two fitness components (nodulation competitiveness for host entry and proliferation), computed for three phenotypic dimensions. **Figure 6 – figure supplement 3:** Representative distributions of fitness effects of new mutations for three levels of pleiotropy between nodulation competitiveness and within-host proliferation. **Figure 6 – figure supplement 4:** Effect of the nodulation bottleneck on the relative strength of selection for nodulation competitiveness and proliferation (evolutionary parameters: k = l = 10 and m = 0). **Figure 6 – figure supplement 5:** Effect of the nodulation bottleneck on the relative strength of selection for nodulation competitiveness and proliferation, with a higher probability of beneficial mutations (evolutionary parameters: k = l = 3 and m = 0). **Figure 6 – figure supplement 6:** Effect of the nodulation bottleneck on the relative strength of selection for nodulation competitiveness and proliferation, with a lower probability of beneficial mutations (evolutionary parameters: k = l = 20 and m = 0). **Figure 6 – figure supplement 7:** Effect of the fitness of the ancestor on the relative strength of selection for nodulation competitiveness and proliferation (evolutionary parameters: k = l = 10 and m = 0). **Figure 6 – figure supplement 8:** Effect of the fitness of the ancestor on the relative strength of selection for nodulation competitiveness and proliferation, with a higher probability of beneficial mutations (evolutionary parameters: k = l = 3 and m = 0). **Figure 6 – figure supplement 9:** Effect of the fitness of the ancestor on the relative strength of selection for nodulation competitiveness and proliferation, with a lower probability of beneficial mutations (evolutionary parameters: k = l = 20 and m = 0). **Figure 6 – figure supplement 10:** Effect of the nodulation bottleneck on the relative strength of selection for nodulation competitiveness and proliferation under weak partial pleiotropy (evolutionary parameters: k = l = 6 and m = 4). **Figure 6 – figure supplement 11:** Effect of the nodulation bottleneck on the relative strength of selection for nodulation competitiveness and proliferation under strong partial pleiotropy (evolutionary parameters: k = l = 2 and m = 8). **Figure 6 – figure supplement 12:** Effect of the chronology of symbiotic events and the size of nodulation bottleneck on the relative strength of selection for nodulation competitiveness and proliferation (evolutionary parameters: k = l = 10 and m = 0). **Figure 6 – source data 1:** Raw data from evolutionary simulations are available at: https://doi.org/10.15454/QYB2S9

In our experiment, the ancestral nodulating strains had a much higher potential for improvement in nodulation competitiveness relative to proliferation (5×10^4^ vs. 10^2^–10^3^ fold, respectively, compared to the natural symbiont *C. taiwanensis*) which could explain the faster evolution of the former trait. We performed additional simulations to test how evolutionary trajectories are affected by the genotype of the ancestor. As expected, starting from a higher proliferation fitness in the ancestor further increased the overall fold effect of early selected mutations on nodulation competitiveness (Figure 6c and d). In contrast, starting with a higher initial nodulation competitiveness value of 10^−3^ or 10^−2^ in the ancestor was not sufficient to lead to a stronger selection for proliferation, further confirming the asymmetry between selective pressures acting on the two phenotypes. These findings were qualitatively robust to our assumptions regarding the genetic architecture of the two symbiotic phenotypes (Supplementary file 8, Figure 6 – figure supplement 2–11).

Finally, we wondered if the chronology of symbiotic steps impacts on selective pressures and evolutionary trajectories. We modified the simulations so that clonal multiplication of bacteria (according to their proliferation fitness) occurs before the selective bottleneck. Although this model is not relevant in the context of the legume-rhizobium symbiosis, it may apply to other symbiotic or pathogenic life cycles, for example when a selective bottleneck occurs during dissemination in the environment or within the host^31,32^, as well as in the case of vector-mediated transmitted pathogens^33,34^ or vertically transmitted symbionts^35^. In this context, the dominance of nodulation competitiveness was strongly reduced, bringing the two selective forces close to equilibrium (Figure 6e and f and Figure 6 – figure supplement 12), in agreement with the expectation that each fitness component should have a similar influence on adaptation when they have a multiplicative effect on global fitness. Therefore, the dominance of nodulation competitiveness in the previous simulation runs is likely a consequence of the selective bottleneck occurring before within-host bacterial multiplication and just after the transient hypermutagenesis phase in the rhizosphere.

Altogether, our simulations show that the selective pressures that are experienced by symbiotic bacteria are asymmetric, with a dominance of selection for nodulation competitiveness over within-host proliferation. This asymmetry occurs when a highly diverse bacterial population from the rhizosphere is exposed to the strong selective bottleneck at host entry, allowing the efficient selection of the most competitive clones by the host plant. The fact that the ancestor from our evolution experiment has a lower nodulation competitiveness value compared to proliferation (relative to that of the wild-type rhizobium *C. taiwanensis*) is an additional factor expected to have contributed to the stronger selection on nodulation competitiveness. Importantly, the key factors responsible for the dominance of selection for host entry competitiveness in our model (strong bottleneck at host entry and phenotypic diversity of environmental microbial populations) are found in many symbiotic systems, suggesting that competition for host-entry is probably a widespread driver of bacterial adaptation in diverse symbiotic associations.

## Discussion

Horizontally transmitted symbiotic bacteria have complex lifecycles during which they have to face multiple environmental constraints, both outside and inside the host. Assessing the relative influence of each of these selective pressures on the adaptive trajectories of emerging symbiotic bacteria is critically needed to better understand the eco-evolutionary dynamics of microbial populations^5^.

Here we show that selection on competitiveness for host entry is the dominant selective pressure shaping early adaptation of new rhizobia. This trait, mainly mediated by the plant, improved very fast during our evolution experiment, to a level comparable to that of the natural *Mimosa* symbiont *C. taiwanensis*, and all adaptive mutations identified in lineages B and G improved nodulation competitiveness. Evolutionary simulations led us to propose that the dominance of the selection on host entry over the selection on within-host proliferation that we observed in our system is strongly dependent on the strength of the selective bottleneck that occurs at host entry and its position the life cycle of symbionts. These results shed light on the importance of selective bottlenecks in shaping the evolutionary trajectories of emerging microbial symbionts. Most theoretical^36,37^ and experimental^38–43^ works on the influence of bottlenecks on microbial adaptation have focused on non-selective bottlenecks that randomly purge genetic diversity and reduce the efficiency of natural selection. Yet, another aspect of bottlenecks emerges when considering that transmission and host colonization can be, at least partially, dependent on microbial genotype, prompting us to consider infection bottlenecks as selective events^44^. Selective bottlenecks have already been described during virus infections^31,45–48^ but received less attention in bacteria, except for their role in the evolution of phenotypic heterogeneity^17,49–51^. Yet, potentially all symbiotic bacterial populations (whether commensal, pathogenic or mutualistic) experience bottlenecks of various intensities (sometimes going down to only a few cells^52^) during their life cycles^27,53,54^. Since competitiveness for host entry is a complex, polygenic phenotypic trait^7,55–58^ that often displays extensive variation within natural populations^59,60^, bottlenecks are expected to represent major selective events (instead of purely stochastic events) in many symbiotic life cycles. Interestingly, strong selection for host entry in experimentally evolved mutualistic and pathogenic symbionts was recently observed in several studies^61,62^ reinforcing the idea that selective bottlenecks at host entry may play a crucial role in the evolution of diverse host-microbe interactions.

In multistep infection processes, the different infection steps can be functionally linked either by couplings or trade-offs^63,64^. Our study unveiled a new characteristic of the rhizobium-legume interactions: the recurrent genetic coupling between nodulation competitiveness and within-host proliferation. Indeed, all the 10 identified mutations that improved within-host proliferation also improved nodulation competiveness, while the reverse was not true. This result corroborates our previous data showing that mutations promoting nodule cell infection always improved nodule formation^14,65^. It is conceivable that the intracellular release and proliferation of rhizobia in nodule cells is dependent on the efficiency of the earliest symbiotic events, *i. e*. the entry and progression of bacteria in infection threads and the concomitant divisions of nodule cells preparing for the accommodation of bacteria. Consistently, delayed progression of infection threads across the root cell layers was shown to impair the release of bacteria in nodule cells^66^. Moreover the identification of plant receptors (LYK3, NFR1, Sym37, SYMRK) and transcription factors (NIN, ERN1, ERN2) involved in both nodule organogenesis and rhizobial intracellular infection^67–70^ supports the existence of common mechanisms controlling the two processes on the plant side. Future functional analyses of the adaptive mutations identified in this study will expand our understanding of the molecular bases of nodulation competitiveness in rhizobia^60,71,72^, and its relationship to within-host proliferation. Including mutualistic traits (nitrogen fixation and host growth promotion) in the analysis of the genetic couplings (or trade-offs) between the different symbiotic traits will be important to fully characterize the genetic constraints shaping the evolution of rhizobium-legume interactions^72–74^. The genetic links between different host colonization phases are generally poorly documented in other symbiotic systems, but they were analyzed in two recent experimental evolution studies with opposite outcomes. Robinson et al. (2018) showed that early bacterial adaptation to zebrafish gut favored the improvement of host entry and inter-host transmission without affecting within-host proliferation^61^. In another study, experimental evolution of *Vibrio fisherii* in symbiosis with squids led to the fixation of mutations in a global regulatory gene (*binK*) that improved both the initiation and maintenance of the interaction, through its action on several bacterial phenotypic traits^75^. Conversely, an example of trade-off between early and late infection stages was recently evidenced in *Xenorhabdus nematophila* interacting with insects^76,77^. In the same line, a trade-off between within-vector and within-host fitness was evidenced in the mosquito-borne parasite *Plasmodium falciparum* causing malaria^78^. These examples highlight the diversity and context-dependence of genetic correlations that exist between bacterial phenotypic traits involved in symbiotic interactions.

In all lineages of this experiment, the rate of adaptation was very high during the first cycles of evolution and then tended to decrease over time. This classical pattern of evolution, due to diminishing return epistasis among beneficial mutations^24,79,80^, was likely favored by the low initial fitness values of nodulating ancestors compared to the natural symbiont *C. taiwanensis* allowing the rapid acquisition of highly beneficial mutations during the first cycles (Table 1). Elevated mutation rate in the rhizosphere, prior to the entry of bacteria in roots^18^, has also played an important role in the dynamic of adaptation by fueling the extensive genetic diversification of bacteria and exposing these populations to plant-mediated selection. A probable consequence of high mutation rate combined to strong selection is the presence of large cohorts of mutations that carry multiple beneficial mutations along with some neutral or slightly deleterious ones. Mutational cohorts were already described in previous evolution experiments^21,81^, with cases of cohorts carrying 2 co-driver mutations^22^. Here we identified up to 7 adaptive mutations in one of our cohorts. The occurrence of multi-driver mutations per cohort might be explained by the nested, sequential fixation of multiple adaptive mutations in one lineage before it reaches detectable frequency (5% in our case)^22^. This phenomenon, creating a ‘travelling wave’ of adaptation^82^, was recently observed with high-resolution sequencing^23^ and is possibly amplified by the strong selection/high mutation regime of our experiment. Alternatively, it was proposed that, in the case of hypermutagenesis, the likelihood of multiple adaptive mutations arising simultaneously in a given genome becomes non-negligible and might promote saltational evolution^83^.

In conclusion, our work shows that the relative strengths of the selective forces imposed by hosts strongly influence the evolutionary trajectory of microbial populations. Selection for host entry is likely the predominant life-history trait selected during the evolution of new horizontally-transmitted symbiotic interactions, such as the rhizobium-legume symbiosis. This effect emerges from the presence of a selective bottleneck at host entry, although its predominance can be modulated by other genetic and ecological factors. Investigating the role of selective bottlenecks in other symbiotic interactions will improve our ability to manage and predict the eco-evolutionary dynamics of host-associated microbial populations.

## Methods

### Bacterial strains and growth conditions

Strains and plasmids used in this study are listed in the Key ressources table. *Cupriavidus taiwanensis* strains were grown at 28°C on tryptone-yeast (TY) medium^84^. *Ralstonia solanacearum* strains were grown at 28°C either on BG medium^85^ or on minimal MP medium^86^ supplemented with 2% glycerol. *Escherichia coli* strains were grown at 37°C on Luria-Bertani medium^87^. Antibiotics were used at the following concentrations: trimethoprim at 100 μg ml^−1^, spectinomycin at 40 μg ml^−1^, streptomycin at 200 μg ml^−1^ and kanamycin at 25 μg ml^−1^ (for *E. coli*) or 50 μg ml^−1^ (for *R. solanacearum*).

### Experimental evolution

Symbiotically evolved clones and populations were generated as previously described^16^. Five lineages, two (B and F) derived from the CBM212 ancestor, two (G and K) derived from the CBM349 ancestor and one (M) derived from the CBM356 ancestor, previously evolved for 16 cycles were further evolved until cycle 35. For each lineage, at each cycle, 30 plants were inoculated with nodule bacterial populations from the previous cycle. Twenty-one days after inoculation, all nodules from the 30 plants were pooled, surface-sterilized with 2.6% sodium hypochlorite for 15 min, rinsed with sterile water and crushed. Then, 10% of the nodule crush was used to inoculate a new set of 30 plants the same day. Serial dilutions of each nodule crush were plated and one clone was randomly selected from the highest dilution and purified. The selected clones, the rest of the nodule crushes, and a 24-h culture of bacteria from an aliquot of nodule crushes were stored at −80°C at each cycle.

### Sequencing bacterial populations and clones

Aliquots of frozen nodule bacterial populations or single colonies from purified clones were grown overnight in BG medium supplemented with trimethoprim. Bacterial DNA was extracted from 1 ml of culture using the Wizard genomic DNA purification kit (Promega). Evolved population DNAs were sequenced at the GeT-PlaGe core facility (https://get.genotoul.fr/), INRAE Toulouse. DNA-seq libraries have been prepared according to Illumina’s protocols using the Illumina TruSeq Nano DNA HT Library Prep Kit. Briefly, DNA was fragmented by sonication, size selection was performed using SPB beads (kit beads) and adaptators were ligated to be sequenced. Library quality was assessed using an Advanced Analytical Fragment Analyzer and libraries were quantified by qPCR using the Kapa Library Quantification Kit (Roche). Sequencing has been performed on a NovaSeq6000 S4 lane (Illumina, California, USA) using a paired-end read length of 2×150 pb with the Illumina NovaSeq Reagent Kits.

Evolved clones were re-sequenced either by C.E.A/IG/Genoscope using the Illumina GA2X technology (clones B8, B16, F16, G8, G16, K16 and M16), or by the GeT_PlaGe core facility using the Illumina technology HiSeq2000 (clones B1, B2, B3, B4, B5, B6, B7, B9, B10, B11, B12, B13, B14, B15, G1, G2, G3, G4, G5, G6, G7, G9, G10, G11, G12, G13, G14, and G15), or HiSeq3000 (clones B20, B25, B30, G25 and G30), or NovaSeq6000 (clones B35, F35, G35, K35 and M35).

### Detection of mutations and molecular analyses

Sequencing reads from NovaSeq6000 runs (whole-populations and clones from cycle 35) were mapped on the chimeric reference genome of the ancestral strain, comprising *R. solanacearum* GMI1000 chromosome (GenBank accession number: NC_003295.1) and megaplasmid (NC_003296.1) together with *C. taiwanensis* symbiotic plasmid pRalta (CU633751). Mutations were detected using breseq v0.33.1(Ref.^88^) with default parameters, either using the polymorphism mode (for whole-population sequences) or the consensus mode (for individual clones). Mutation lists were curated manually in order to remove mutations present in the ancestral strains as well as false positive hits arising from reads misalignments in low complexity regions. Moreover, alleles showing aberrant trajectories in the time-course whole-population sequencing data were checked manually, by inspecting either breseq output files and/or read alignments with IGV^89^, and corrected as needed. In this work, we focused our attention on SNPs and indels detected above 5% frequency in the populations. Recombinations, rearrangements and IS movements were not analyzed exhaustively. Mutations detected in populations are listed in Figure 2 – source data 1.

Sequencing data of the remaining clones were analyzed as described previously^26^ with the PALOMA bioinformatics pipeline implemented in the Microscope platform^90^. The complete list of mutational events generated for these clones are available on the Microscope platform (https://mage.genoscope.cns.fr/microscope/expdata/NGSProjectEvo.php, SYMPA tag). Mutations detected in clones are listed in Figure 2 – source data 2.

In order to identify mutations forming temporal clusters (cohorts) through populations, we selected mutations having a frequency of 30% in at least one cycle and clustered their frequency using the *hclust* function in R v3.6.1^91^. Then, we separated cohorts using a cutoff distance of 0.3.

Sub-population genealogies in lineages B and G, shown as Muller plots in Figure 2 – figure supplement 1, were reconstructed by comparing mutations found in cohorts and in individually sequenced clones. Cohorts for which ancestry cannot be ascertained (mutations not found in any clone) were not included in these plots. Relative frequencies of genotypes were calculated manually and plotted with the R package MullerPlot^92^.

G scores were calculated as described^20^. Since we found that synonymous mutations can be adaptive, we used all types of mutations in coding regions to calculate this statistics. Moreover, we included all mutations beyond 5% frequency in this analysis since we assumed that strong clonal interference may prevent adaptive mutations to rise to high frequency. To evaluate statistical significance of G scores, we ran 1,000 simulations where the total number of mutations used for the calculation of the observed G scores (3330) were randomly attributed in the coding genes according to their respective length to compute simulated G scores for each bacterial gene. The sum of simulated G scores was compared to the observed sum and we computed a Z score and *P*-value from these simulated G statistics. These simulations were also used to compute mean G scores for each gene, and to calculate the associated Z scores and *P*-values (adjusted using a Bonferroni correction).

### Reconstruction of mutations in evolved clones

Mutant alleles and constitutively expressed reporter genes (GFPuv, mCherry) were introduced into *Ralstonia* evolved clones by co-transformation using the MuGent technique^26,93^. Briefly, two DNA fragments, the first one carrying an antibiotic (kanamycin) resistance gene prepared from the pRCK-P*ps*-GFP or pRCK-P*ps*-mCherry linearized plasmids allowing the integration of the resistance gene in the intergenic region downstream the *glmS* gene, and the second one carrying the mutation to be introduced prepared by PCR amplification of a 6 kb region using genomic DNA of evolved clones as template and high fidelity Phusion polymerase (ThermoFisher Scientific), were co-transformed into naturally competents cells of *Ralstonia* evolved clones. Co-transformants resistant to kanamycin were screened by PCR using primers specifically amplifying mutant or wild-type alleles and verified by Sanger sequencing. Primers used in mutation reconstructions are listed in the Key ressources table.

### Analyses of symbiotic phenotypes on *M. pudica*

*Mimosa pudica* seeds (LIPME 2019 production obtained from one commercial seed (B&T World Seed, Paguignan, France) of Australian origin) were sterilized as described^94^ by immersion in 95% H2SO4 during 15 minutes and 2.4% sodium hypochlorite solution during 10 minutes and rinsed in sterile distilled water 5 times. Seeds were soaked in sterile water at 28°C under agitation for 24 hours and then deposited on soft agar (9.375 g/L) and incubated at 28°C during 24 more hours in darkness. Then, seedlings were cultivated in glass tubes (2 seedlings/tube) under N-free conditions, each tube containing 20 mL of solid Fahraeus medium^95^ and 40 mL of liquid Jensen medium^96^ diluted 1:4 with sterile water. Plants were incubated in a culture chamber at 28°C under a photoperiod day/night of 16 h/8 h and positioned randomly in the different experiments.

For *in planta* relative fitness and nodulation competitiveness assays, two strains of *Ralstonia solanacearum* expressing differential constitutive fluorophores (GFPuv or mCherry) or one strain of *R. solanacearum* and one strain of *C. taiwanensis*, both expressing differential antibiotic resistance (streptomycin and trimethoprim), were co-inoculated onto *M. pudica* plantlets grown for 3 to 4 days in the culture chamber. Both strains were inoculated in equivalent proportion (*ca*. 5×10^5^ bacteria of each strain per plant) except for the comparisons of the *Ralstonia* nodulating ancestors with *C. taiwanensis* where strains were mixed in a 1000:1 ratio (*ca*. 5×10^3^ bacteria of *C. taiwanensis* and *ca*. 5×10^5^ bacteria of nodulating ancestors per plant). Nodules were harvested 21 days after inoculation, surface sterilized by immersion in 2.4% sodium hypochlorite solution for 15 minutes and rinsed with sterile water. For *in planta* relative fitness measurements, sterilized nodules from 20 plants (see the exact number of harvested nodules in Supplementary file 1) were pooled and crushed in 1 mL of sterile water, diluted and spread on selective solid medium using an easySpiral automatic plater (Interscience). After 2-day incubation at 28°C, colonies were screened either by plating on selective medium in case of co-inoculations of *Ralstonia* evolved clones with *C. taiwanensis* or based on fluorescence using a stereo zoom microscope (AxioZoom V16, Zeiss, Oberkochen, Germany) for co-inoculations of *Ralstonia* evolved clones with *Ralstonia* reconstructed mutants. For nodulation competitiveness assays, *ca*. 96 individual nodules per experiment were crushed separately in 96-well microtiter plates and droplets were deposited on selective medium. Nodule occupancy was determined by screening bacteria grown in the droplets either on selective medium or based on their fluorescence as described for fitness measurements. Both nodulation competitiveness and relative fitness assays were measured in at least three independent biological replicates. Competitive indexes (CI) were calculated by dividing the ratio of the number of test strain (evolved strains or reconstructed mutants) vs. reference strain (*C. taiwanensis* or evolved parental strains, respectively) in nodules normalized by the inoculum ratio. When CI values were all above 1, CI values were transformed by their inverse and compared to the value 1 using a one-sided Student *t*-test (*P*<0.05).

For within-host proliferation assays, plantlets were inoculated with a single strain (5×10^5^ bacteria per plant). In each experiment, nodules from 6 individual plants were collected separately 21 days after inoculation, surface sterilized for 15 minutes in a 2.4% sodium hypochlorite solution, rinsed with sterile water and crushed in 1 ml of sterile water. Dilutions of nodule crushes were plated on selective solid medium using an easySpiral automatic plater (Interscience). Two days after incubation at 28°C, the number of colonies was counted. Within-host proliferation was estimated as the number of bacteria per nodule. For each strain, 15 to 24 measurements from three independent biological replicates were performed. Pairwise comparisons of proliferation values were compared using a two-sided Wilcoxon rank sum test.

For Jensen culture medium and rhizosphere colonization assays, plantlets were co-inoculated with pair of strains expressing differential constitutive fluorophores (GFPuv or mCherry) in equivalent proportion (5×10^6^ bacteria of each strain per plant). Seven days after inoculation, bacteria present in the culture medium were diluted and plated on selective solid medium using an easySpiral automatic plater (Interscience). Bacteria attached to the roots were resuspended in 4 ml of sterile water by strongly vortexing for 1 minute, diluting and plating on selective medium using an easySpiral automatic plater (Interscience). Two days after incubation at 28°C, colonies were screened for fluorescence using a stereo zoom microscope (AxioZoom V16, Zeiss, Oberkochen, Germany). Competitive indexes (CI) for Jensen medium or rhizosphere colonization were calculated as described above. CI values were compared to the value 1 using a two-sided Wilcoxon rank sum test with Benjamini-Hochberg correction.

Sample sizes were determined based on our previous studies^26,65,97^. In cases where we performed only three independent replicates, each replicate was based on the analysis of a large number of plants or nodules. Independent biological replicates were performed on different days, with different plants and different bacterial cultures.

### Modelling/simulations

Full details on evolutionary simulations and the list and values of parameters used in the simulations are available in Supplementary files 8 and 9, respectively.

## Data availability statement

Sequencing data are available under NCBI SRA BioProject ID PRJNA788708 and SRP353965. Raw experimental data are available as Supplementary Information. Raw data generated from computer simulations are deposited on the Data INRAE dataverse (https://doi.org/10.15454/QYB2S9).

## Code availability statement

Computer code used to analyze genomic data and to perform computer simulations are deposited on the Data INRAE dataverse (https://doi.org/10.15454/QYB2S9).

## Material availability statement

Requests for biological resources should be directed to the corresponding authors P.R. or D.C. *Ralstonia solanacearum* strains will be shared upon fulfilment of appropriate biosecurity procedures for quarantine bacteria.

## Acknowledgements

We are grateful to A. Carlier and F. Roux for careful reading of the manuscript, to O. Tenaillon for helpful discussions on the project, to O. Tenaillon and H. Le Nagard for sharing their genetic algorithm script, and to Ludovic Legrand for archiving sequencing data and their associated metadata. We acknowledge the Genotoul bioinformatics platform Toulouse Occitanie (Bioinfo Genotoul, https://doi.org/10.15454/1.5572369328961167E12) for providing computing and storage resources.

G.G.D.d.M was supported by a fellowship from the French Ministère de l’Enseignement Supérieur, de la Recherche et de l’Innovation (MESRI). P.R. received funding from the EU in the framework of the Marie-Curie FP7 COFUND People Programme, through the award of an AgreenSkills+ fellowship (under grant agreement n°609398) and from the European Union’s Horizon 2020 research and innovation programme under the Marie Skłodowska-Curie grant agreement N° 845838.

This study was supported by the Fédération de Recherche Agrobiosciences, Interactions et Biodiversité, the French National Research Agency (ANR-16-CE20-0011-01 and ANR-21-CE02-0019-01) and the “Laboratoires d’Excellence (LABEX)” TULIP (ANR-10-LABX-41)” and the “École Universitaire de Recherche (EUR)” TULIP-GS (ANR-18-EURE-0019).

## Authors contributions

C.M.-B, D.C. and P.R. designed the project. G.G.D.M. performed most experiments and analyze the data. S.M., N.G., A.-C.C., D.C. and P.R. performed experiments. J.-B.F. and P.R. performed computer simulations. M.M., T.B. and S.V. performed NGS sequencing at GeT-PlaGe. D.R. analyzed mutations in clones and deposit data in the microScope platform. D.C. and P.R. supervised experimental work and data analysis. C.B.-M., J.-B.F and P.R. acquired funding for the project. G.G.D.M., J.B-F., D.C. and P.R. wrote the manuscript, which was approved by all authors.

## Competing interests

The authors declare no competing interests.

## Supplementary information

### Supplementary files

**Supplementary file 1:** Number of nodules harvested at each evolution cycle.

**Supplementary file 2:** Coverage depth for whole-population sequencing

**Supplementary file 3:** Summary of detected mutations in the five lineages

**Supplementary file 4:** Analysis of genetic parallelism in the five lineages.

**Supplementary file 5:** Global analysis of mutational cohorts

**Supplementary file 6:** Neutral and deleterious reconstructed mutations

**Supplementary file 7:** Frequencies of usage of codons modified by adaptive synonymous mutations

**Supplementary file 8:** Supplementary information on the evolutionary simulations

**Supplementary file 9:** Parameter values used in computer simulations

### Source Data

**Figure 1 – source data 1**. Raw data obtained for the phenotypic characterization of *Ralstonia* evolved clones compared to *C. taiwanensis.*

**Figure 1 – figure supplement 2 – source data 1:** Raw data obtained for the measurements of the dry weights of M. pudica plants inoculated with cycle 35 clones.

**Figure 2 – source data 1.** Mutations identified in the evolved populations from all lineages

**Figure 2 – source data 2:** Mutations present in the evolved clones

**Figure 3 – source data 1:** Raw data obtained for the characterization of the *in planta* fitness of evolved clones carrying fixed mutations.

**Figure 4 – source data 1:** Raw data obtained for the characterization of nodulation competitiveness of evolved clones carrying fixed mutations.

**Figure 4 – figure supplement 1 – source data 1:** Raw data obtained for the Jensen culture medium colonization and rhizosphere colonization assays.

**Figure 5 – source data 1**: Raw data obtained for the characterization of the capacity of evolved clones carrying fixed mutations to proliferate within the host.

**Figure 6 – source data 1:** Raw data from evolutionary simulations are available at: https://doi.org/10.15454/QYB2S9

**Key ressources table:** Strains, plasmids, oligonucleotides and software used and sequence data generated in this study.

### Figure supplements

**Figure 1 – figure supplement 1:** Experimental evolution of *Ralstonia solanacearum* GMI1000 pRalta through serial cycles of plant (*Mimosa pudica*) inoculation-isolation of nodule bacteria.

**Figure 1 – figure supplement 2:** Dry weights of *M. pudica* plants inoculated with cycle 35 evolved clones.

**Figure 2 – figure supplement 1:** Evolution of population composition over time in lines B and G

**Figure 4 – figure supplement 1:** Survival of reconstructed mutants from lineages B and G in the Jensen medium and rhizosphere.

**Figure 6 – figure supplement 1:** Schematic representation of the modelling framework.

**Figure 6 – figure supplement 2**: Distribution of fitness effects of new mutations for each of the two fitness components (nodulation competitivity and proliferation), computed for three phenotypic dimensions.

**Figure 6 – figure supplement 3:** Representative distributions of fitness effects of new mutations for three levels of pleiotropy between nodulation competitiveness and within-host proliferation.

**Figure 6 – figure supplement 4:** Effect of the nodulation bottleneck on the relative strength of selection for nodulation competitiveness and proliferation (evolutionary parameters: k = l = 10 and m = 0).

**Figure 6 – figure supplement 5:** Effect of the nodulation bottleneck on the relative strength of selection for nodulation competitiveness and proliferation, with a higher probability of beneficial mutations (evolutionary parameters: k = l = 3 and m = 0).

**Figure 6 – figure supplement 6:** Effect of the nodulation bottleneck on the relative strength of selection for nodulation competitiveness and proliferation, with a lower probability of beneficial mutations (evolutionary parameters: k = l = 20 and m = 0).

**Figure 6 – figure supplement 7:** Effect of the fitness of the ancestor on the relative strength of selection for nodulation competitiveness and proliferation (evolutionary parameters: k = l = 10 and m = 0).

**Figure 6 – figure supplement 8:** Effect of the fitness of the ancestor on the relative strength of selection for nodulation competitiveness and proliferation, with a higher probability of beneficial mutations (evolutionary parameters: k = l = 3 and m = 0).

**Figure 6 – figure supplement 9:** Effect of the fitness of the ancestor on the relative strength of selection for nodulation competitiveness and proliferation, with a lower probability of beneficial mutations (evolutionary parameters: k = l = 20 and m = 0).

**Figure 6 – figure supplement 10:** Effect of the nodulation bottleneck on the relative strength of selection for nodulation competitiveness and proliferation under weak partial pleiotropy (evolutionary parameters: k = l = 6 and m = 4).

**Figure 6 – figure supplement 11:** Effect of the nodulation bottleneck on the relative strength of selection for nodulation competitiveness and proliferation under strong partial pleiotropy (evolutionary parameters: k = l = 2 and m = 8).

**Figure 6 – figure supplement 12:** Effect of the chronology of symbiotic events and the size of nodulation bottleneck on the relative strength of selection for nodulation competitiveness and proliferation (evolutionary parameters: k = l = 10 and m = 0).

## Notes

### Competing Interest Statement

The authors have declared no competing interest.

