## Supplementary Information for "A selective bottleneck during host entry drives the evolution of new legume symbionts"

### Figure supplements

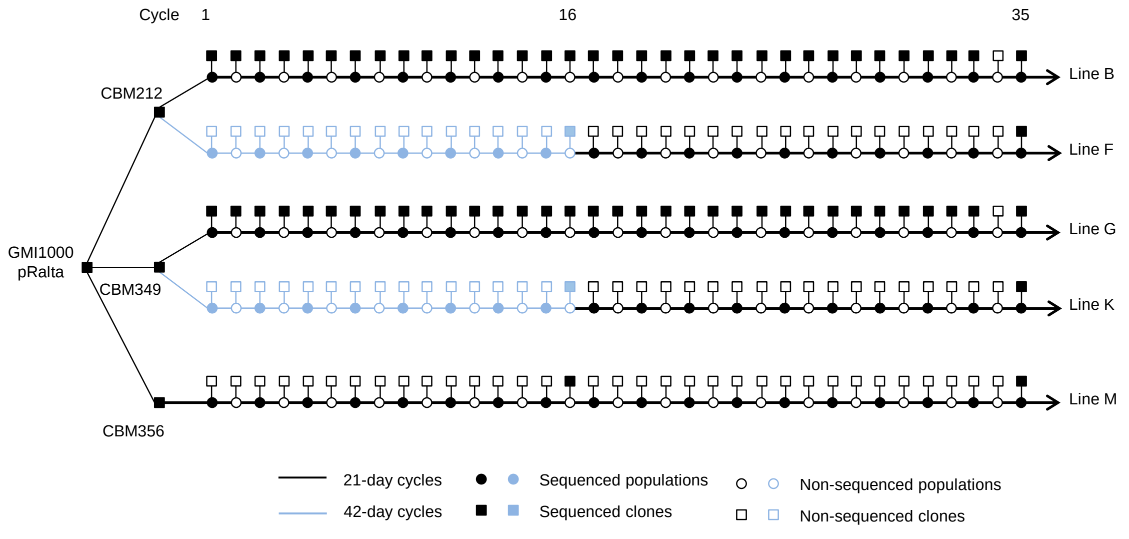


**Figure 1 – figure supplement 1: Experimental evolution of *Ralstonia solanacearum* GMI1000 pRalta through serial cycles of plant (*Mimosa pudica*) inoculation-isolation of nodule bacteria.** The 5 bacterial lineages, lines B, F, G, K, and M were independently derived from the spontaneous nodulating mutants either CBM212, or CBM349, or CBM356. At each evolution cycle, bacterial nodule populations were isolated either 21 days (black lines) or 42 days (blue lines) after plant inoculation and partly directly re-inoculated to new plants to initiate the next cycle. At each cycle, one individual clone representative of the bacterial population was isolated and stored. Sequenced populations and clones are indicated.

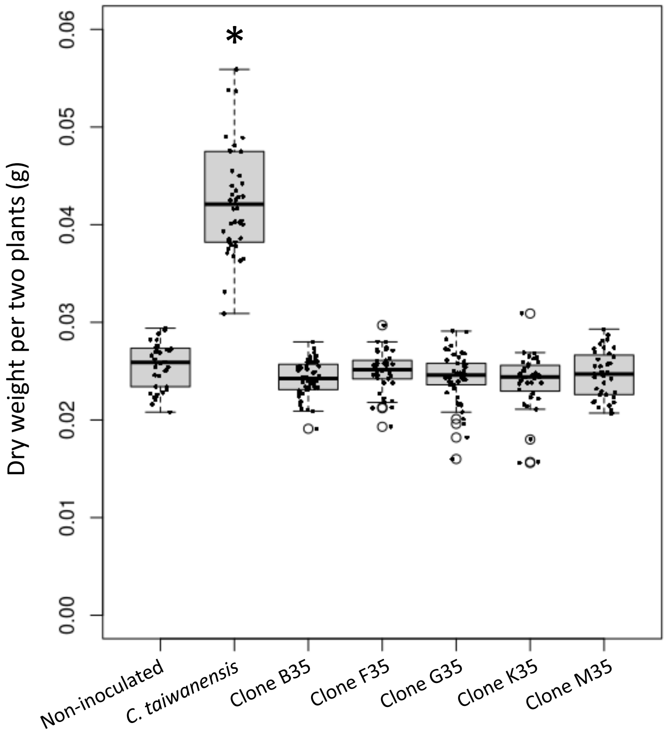


**Figure 1 – figure supplement 2: Dry weights of *M. pudica* plants inoculated with cycle 35 evolved clones.**

The dry weight of the aerial parts of two plants was measured 21 days after inoculation with the evolved clones of cycle 35 or *C. taiwanensis* and compared to non inoculated plants. Data were obtained from 3 to 4 independent experiments for each comparison and the sample size (n) is comprised between n=38-49 . * Significantly different from non inoculated plants (P<0.05, Wilcoxon test).

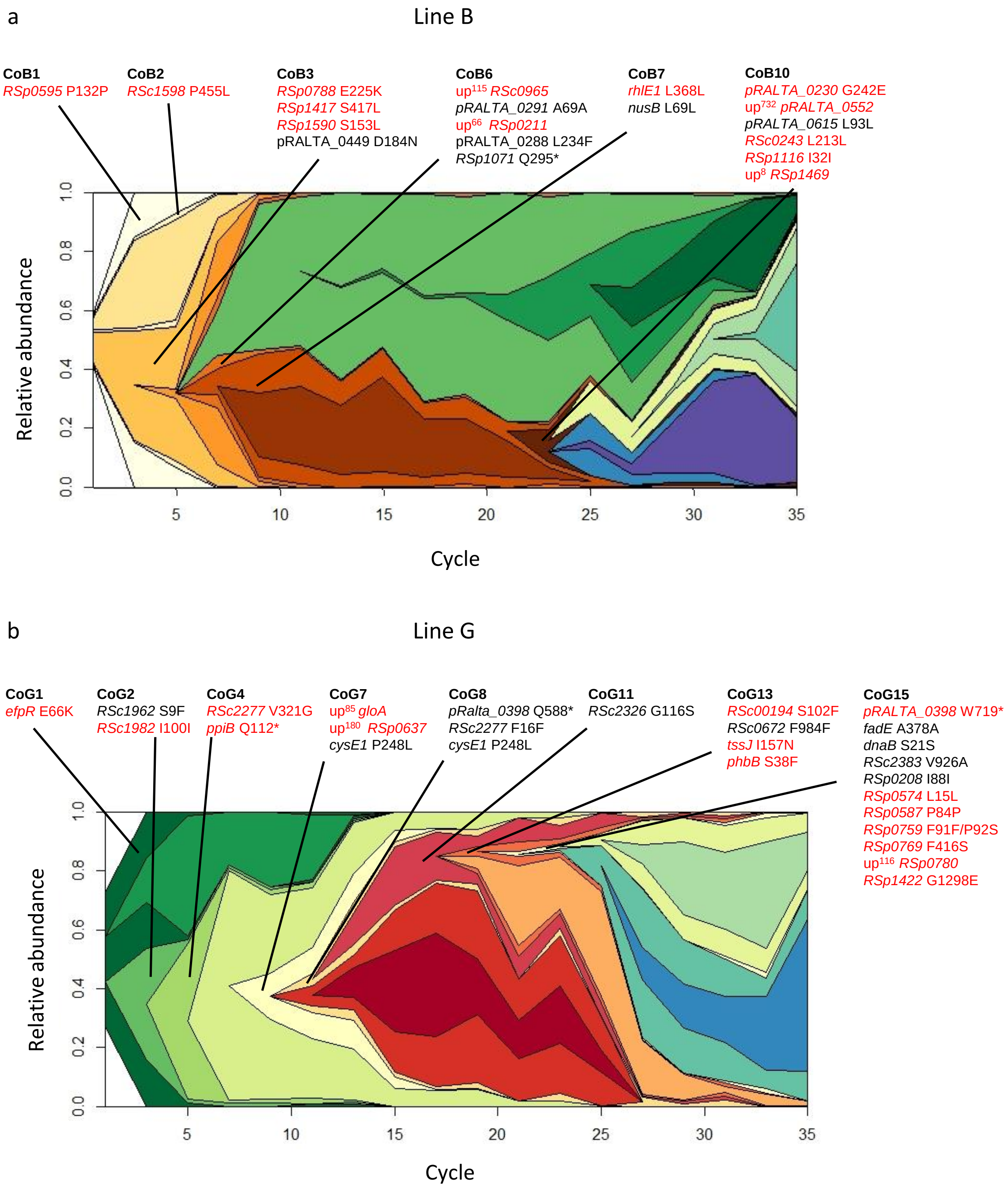


**Figure 2 – figure supplement 1: Evolution of population composition over time in lines B and G.** Muller plots representing the relative frequency of each genotype in lines B (a) and G (b) for each sequenced population (uneven cycles). Each color represent a different genotype, and its relative frequency is shown as the vertical area at each sequenced time point. The genotypes containing cohorts for which adaptive mutations were identified experimentally are indicated. For each cohort, non-adaptive or deleterious mutations are shown in black, adaptive mutations are in red.

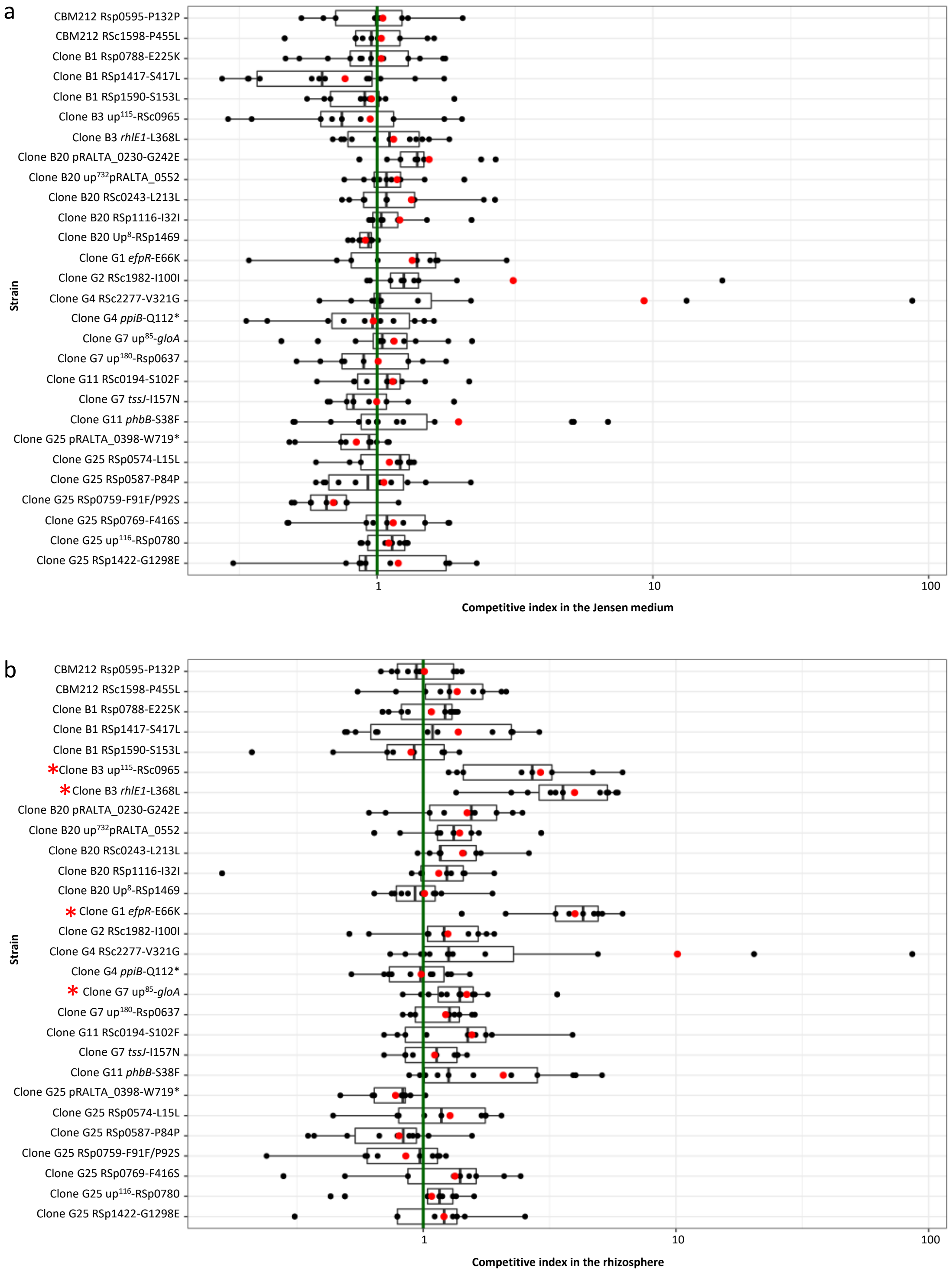


**Figure 4 – figure supplement 1: Survival of reconstructed mutants from lineages B and G in the Jensen medium and rhizosphere**.

Survival of evolved clones carrying reconstructed adaptive mutations from fixed mutational cohorts in the Jensen medium (**a**) and in rhizosphere (**b**). Competitive indexes (CI) were calculated as the ratio of the mutant strain on the isogenic parental strain in Jensen medium populations normalized by the inoculum ratio. Red points correspond to mean values of CI. Rectangles span the first quartiles to the third quartiles, bold segments inside the rectangle show the median. Red stars beside strain names indicate the mutations that significantly improved survival in the rhizosphere compared to the parental evolved clone (*P*<0.05, Wilcoxon test with Benjamini-Hochberg correction). Data were obtained from 3-5 independent experiments for each comparison and the sample size (n) is comprised between n=9-14. Raw data are available in Figure 4 – figure supplement 1 - source data 1.

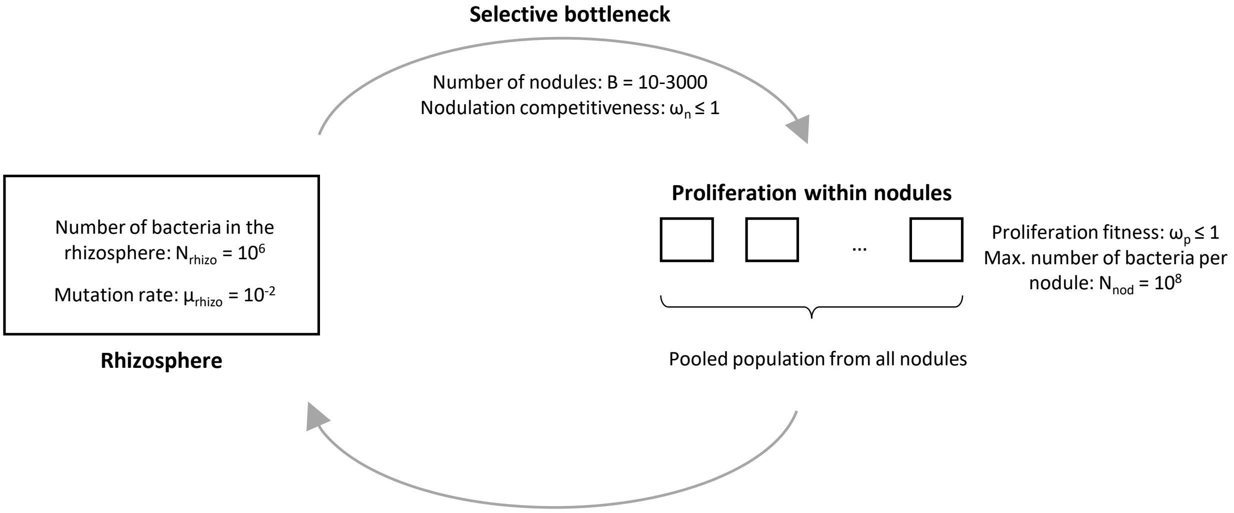


**Figure 6 – figure supplement 1:** Schematic representation of the modelling framework. In the rhizosphere, bacteria acquire mutations (at a rate $\mu_{rhizo}$) that will modify the value of the two fitness components: nodulation competitiveness $\omega_n$) and proliferation capacity within nodules ($\omega_p$). To enter the host, bacteria go through a selective bottleneck, where the probability that each bacterium founds one nodule is equal to its relative nodulation competitiveness value. At each cycle, a fixed number of nodules (B) are formed, defining the stringency of the bottleneck. Once within nodules, bacteria proliferate to a level equal to the product of their proliferation fitness value ($\omega_p$) times the maximal number of bacteria per nodule ($N_{nod}$). At the end of each cycle, all bacteria from all nodules are pooled, and a subset of this population is used to found the rhizospheric population for the next cycle. This step creates a weak non-selective bottleneck with no effect on the evolutionary dynamics since all nodulating clones are represented according to their relative frequency in the nodule population ($B \ll N_{rhizo}$).

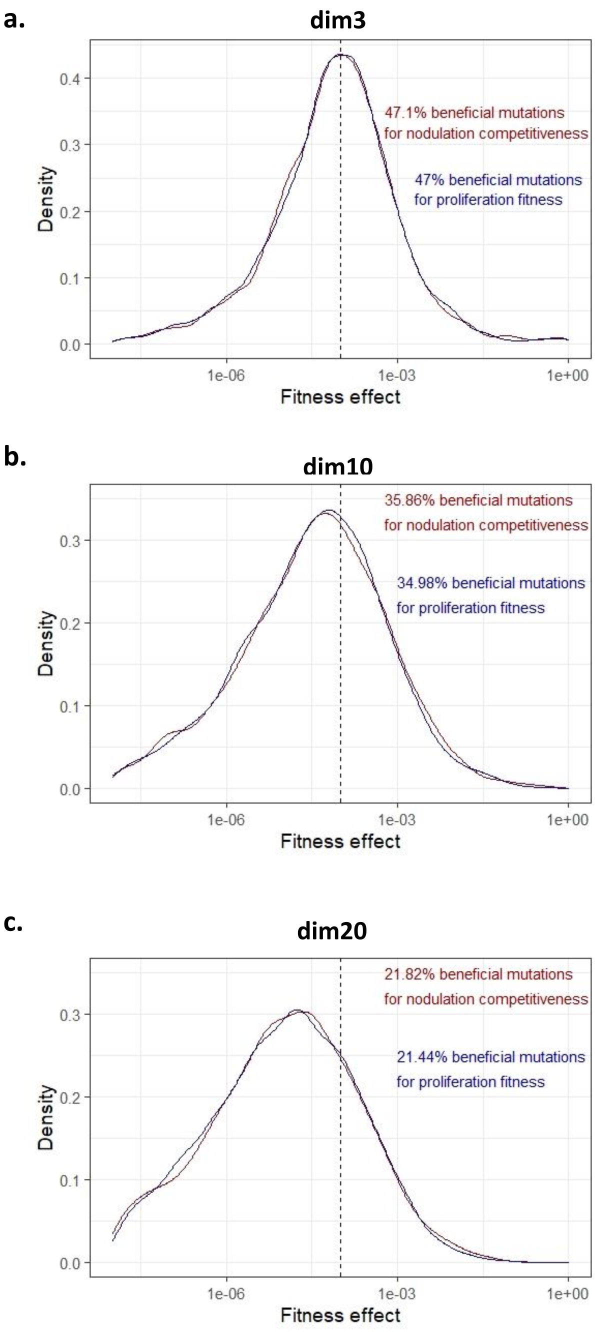


**Figure 6 – figure supplement 2**: Distribution of fitness effects of new mutations for each of the two fitness components (nodulation competitivity and proliferation), computed for three phenotypic dimensions: k = l = 3 (**a**), k = l = 10 (**b**) or k = l = 20 (**c**). In all cases, there is no pleiotropy (m = 0). Distributions were estimated by simulating 1 random mutation in each of 5000 different genetic architectures in the ancestral strain with low fitness ($10^{-4}$ relative to the theoretical optimum for each fitness component, indicated by the vertical dashed line). On each panel, the percentage of beneficial mutations for each component is shown. Differences in the percentage of beneficial mutations between the two fitness components result from the randomness in the definition of the variance-covariance matrices of mutation and fitness effects.

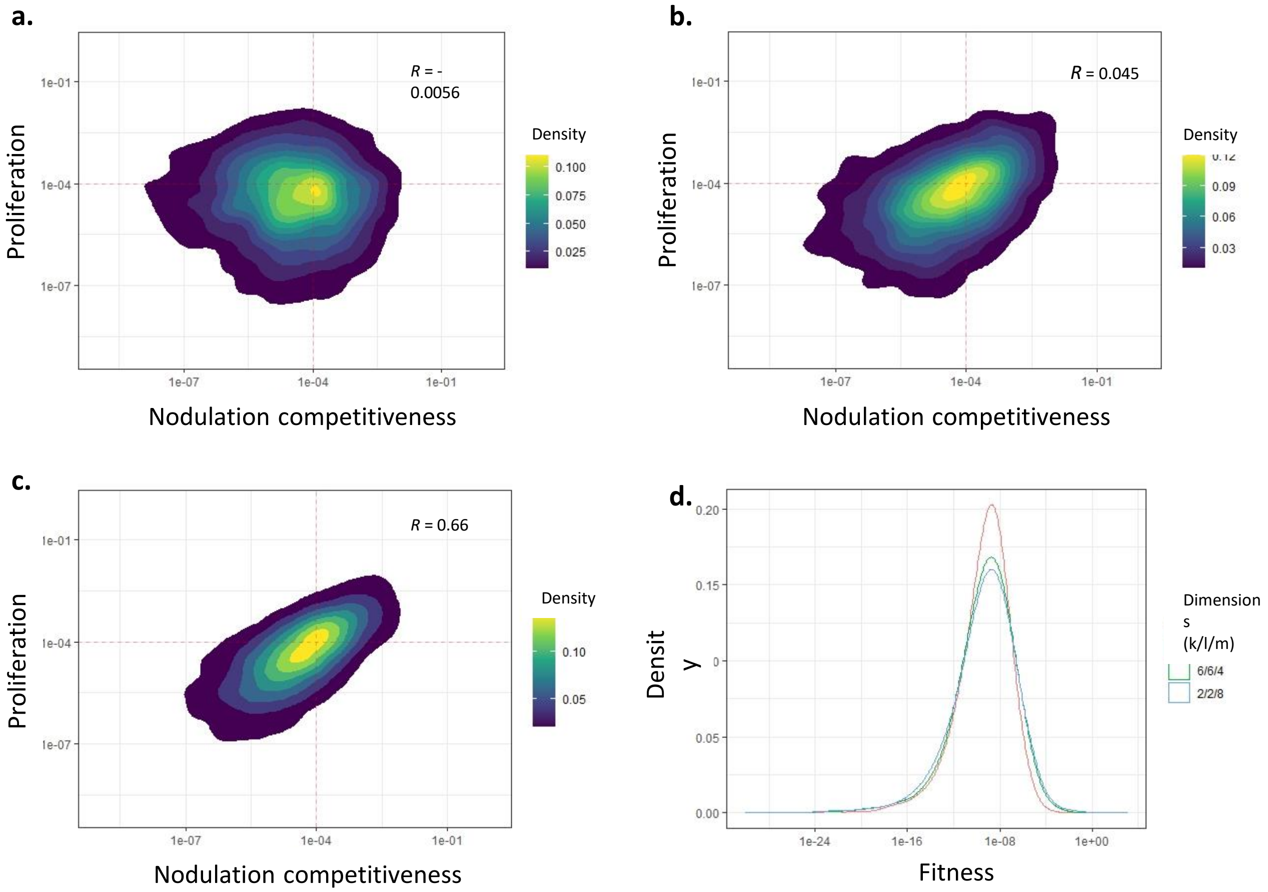


**Figure 6 – figure supplement 3:** Representative distributions of fitness effects of new mutations for three levels of pleiotropy between nodulation competitiveness and within-host proliferation. **a**, No pleiotropy (k = 10, l = 10, m = 0). **b**, Weak partial pleiotropy (k = 6, l = 6, m = 4). **c**, Stronger partial pleiotropy (k = l = 2, m = 8). **d**, Distribution of fitness effects (calculated as the product of nodulation competitiveness by proliferation) for simulated mutations shown in **a-c.** Distributions were computed by simulating 1 mutation in each of 5000 different genetic architectures in the ancestral strain with low fitness ($10^{-4}$ relative to the theoretical optimum for each fitness component, indicated by the red dotted lines).

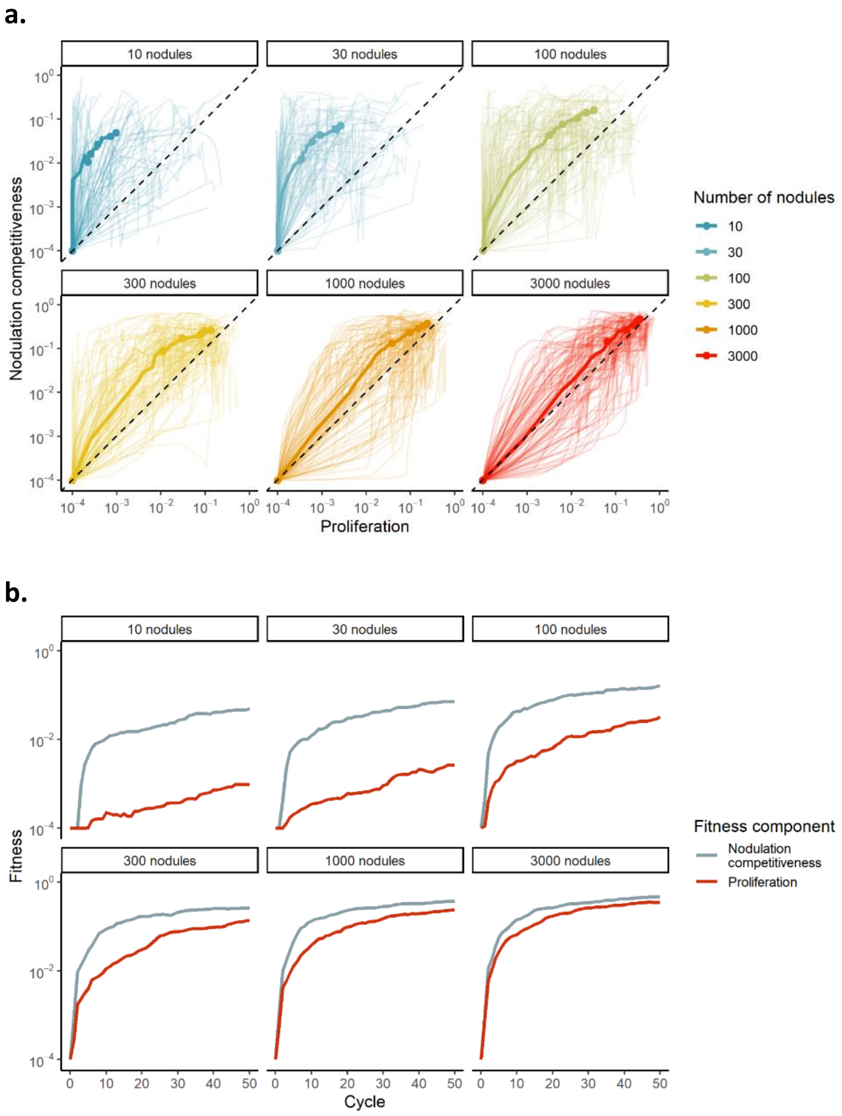


**Figure 6 – figure supplement 4:** Effect of the nodulation bottleneck on the relative strength of selection for nodulation competitiveness and proliferation. Same data as in Figure 4a,b (main text). a, Median (thick lines) and individual (thin lines) fitness trajectories of 100 simulated populations evolving under different sizes of nodulation bottleneck (10 to 3000 nodules). The black dotted line represents the diagonal, along which population would improve both phenotypic traits equally well. b, Improvement of nodulation competitiveness (grey lines) and proliferation (red lines) fitness values along evolutionary cycles under different sizes of nodulation bottlenecks. Evolutionary parameters used were: k = l = 10 and m = 0.

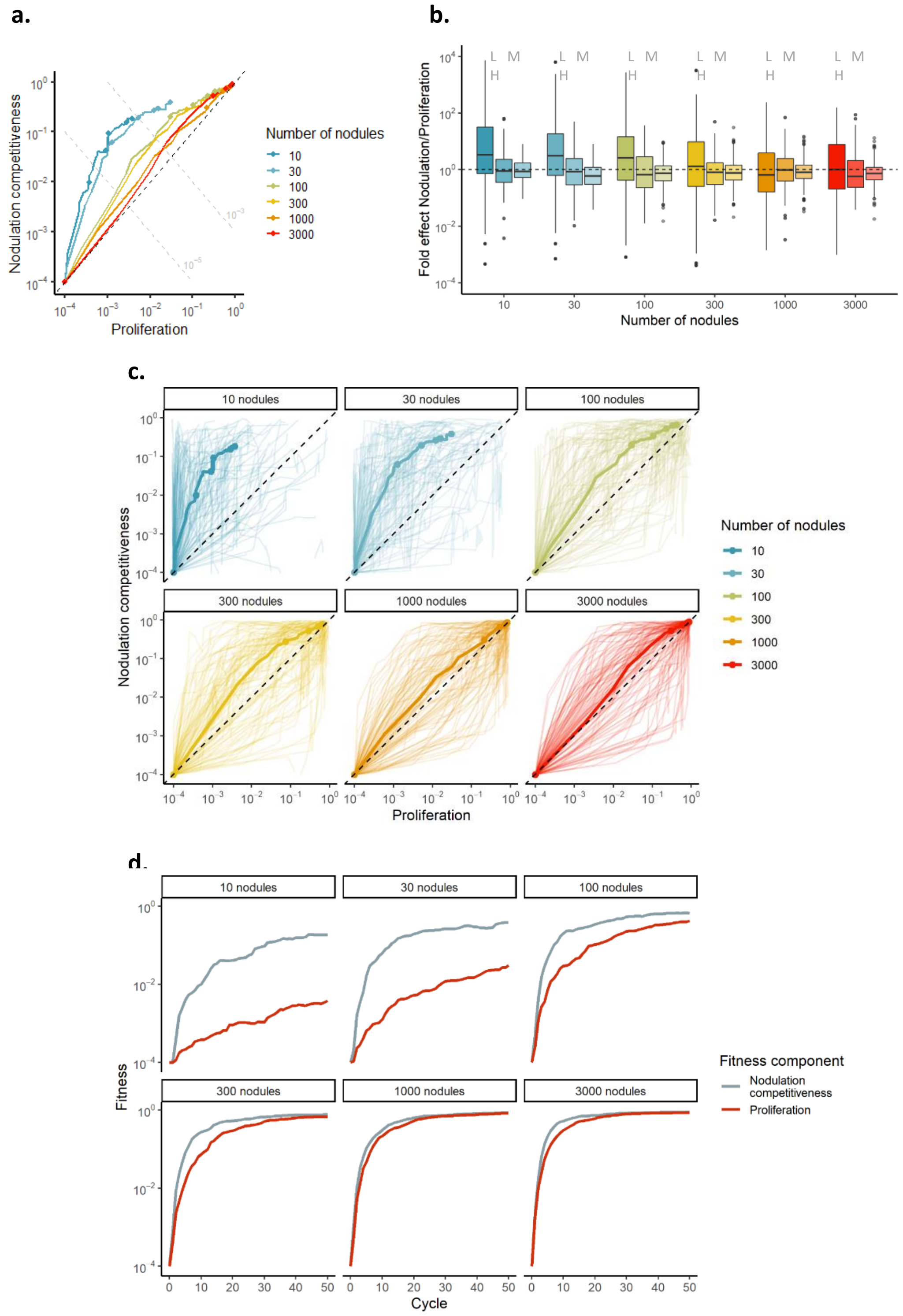


**Figure 6 – figure supplement 5:** Effect of the nodulation bottleneck on the relative strength of selection for nodulation competitiveness and proliferation, with a higher probability of beneficial mutations. a, b, as in Figure 4a,b (main text). c,d, as in Supplementary Figure 4a,b. Evolutionary parameters used were: k = l = 3 and m = 0.

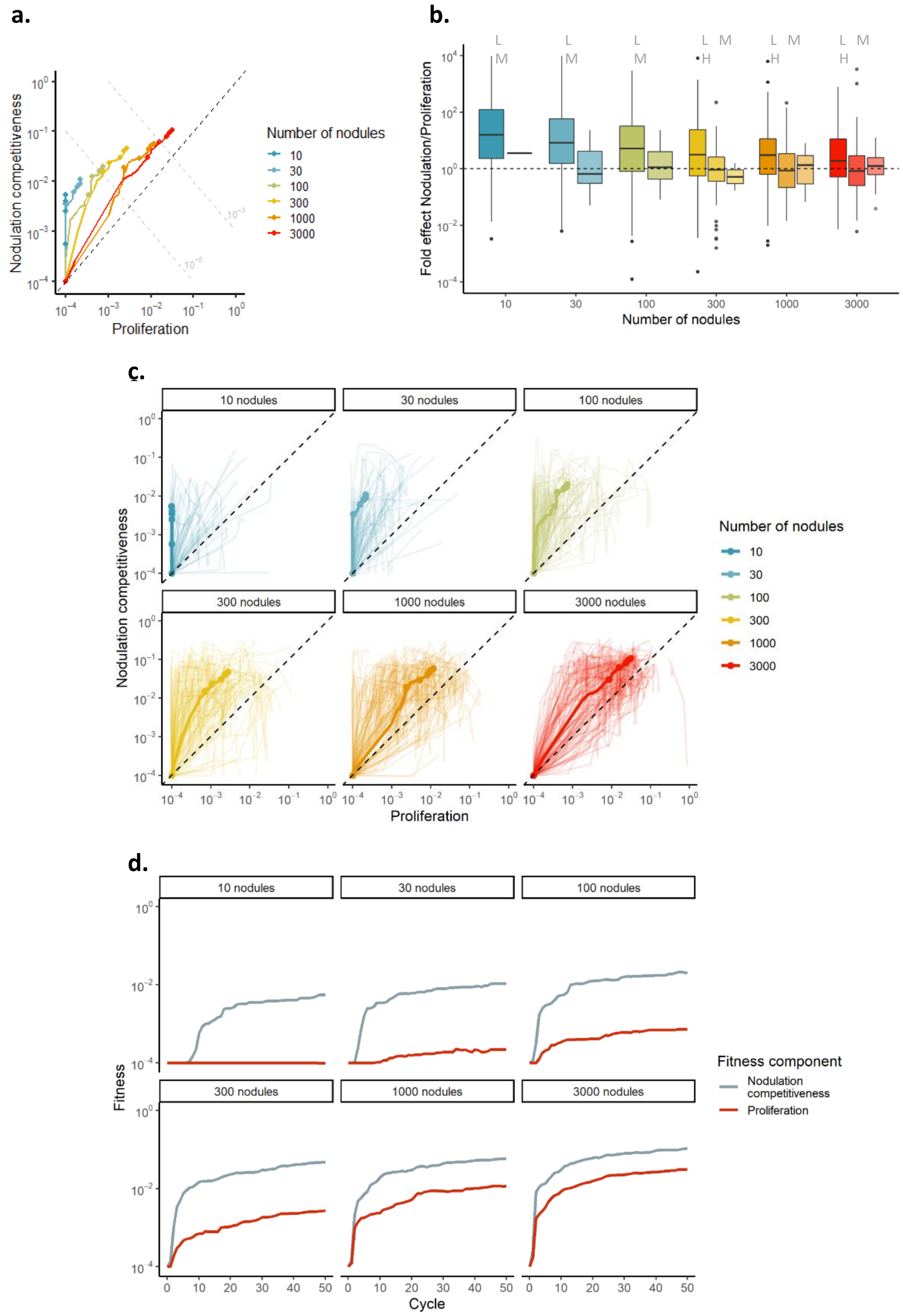


**Figure 6 – figure supplement 6:** Effect of the nodulation bottleneck on the relative strength of selection for nodulation competitiveness and proliferation, with a lower probability of beneficial mutations. a, b, as in Figure 4a,b (main text). c,d, as in Supplementary Figure 4a,b. Evolutionary parameters used were: $k = l = 20$ and $m = 0$.

**a.**

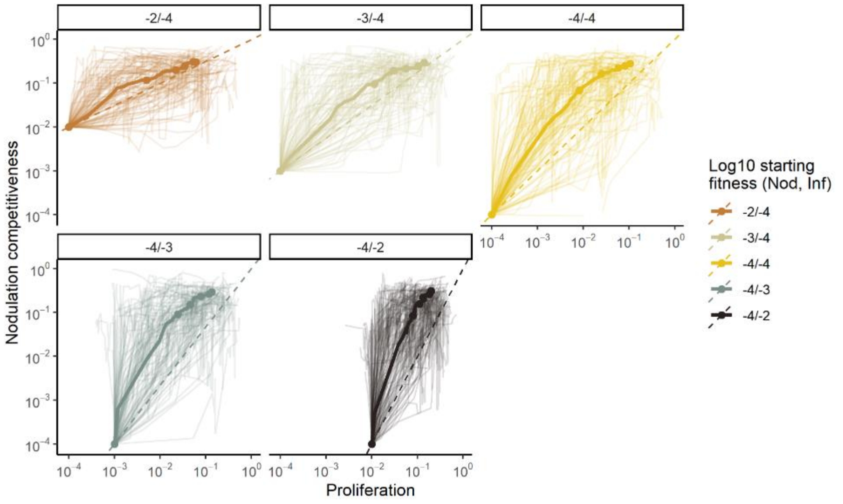


**b.**

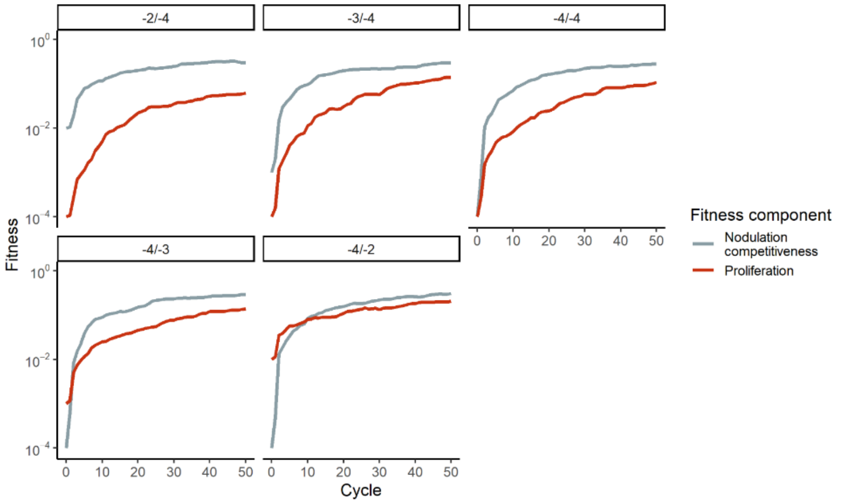


**Figure 6 – figure supplement 7:** Effect of the fitness of the ancestor on the relative strength of selection for nodulation competitiveness and proliferation. Same data as in Figure 4c,d. a, b, as in Supplementary Figure 4a,b, with fitness trajectories shown for various combinations of nodulation competitiveness/proliferations fitness values in the ancestor ($10^{-2}/10^{-4}$), ($10^{-2}/10^{-4}$), ($10^{-4}/10^{-4}$), ($10^{-4}/10^{-3}$), and ($10^{-4}/10^{-2}$). Evolutionary parameters used were: k = l = 10 and m = 0.

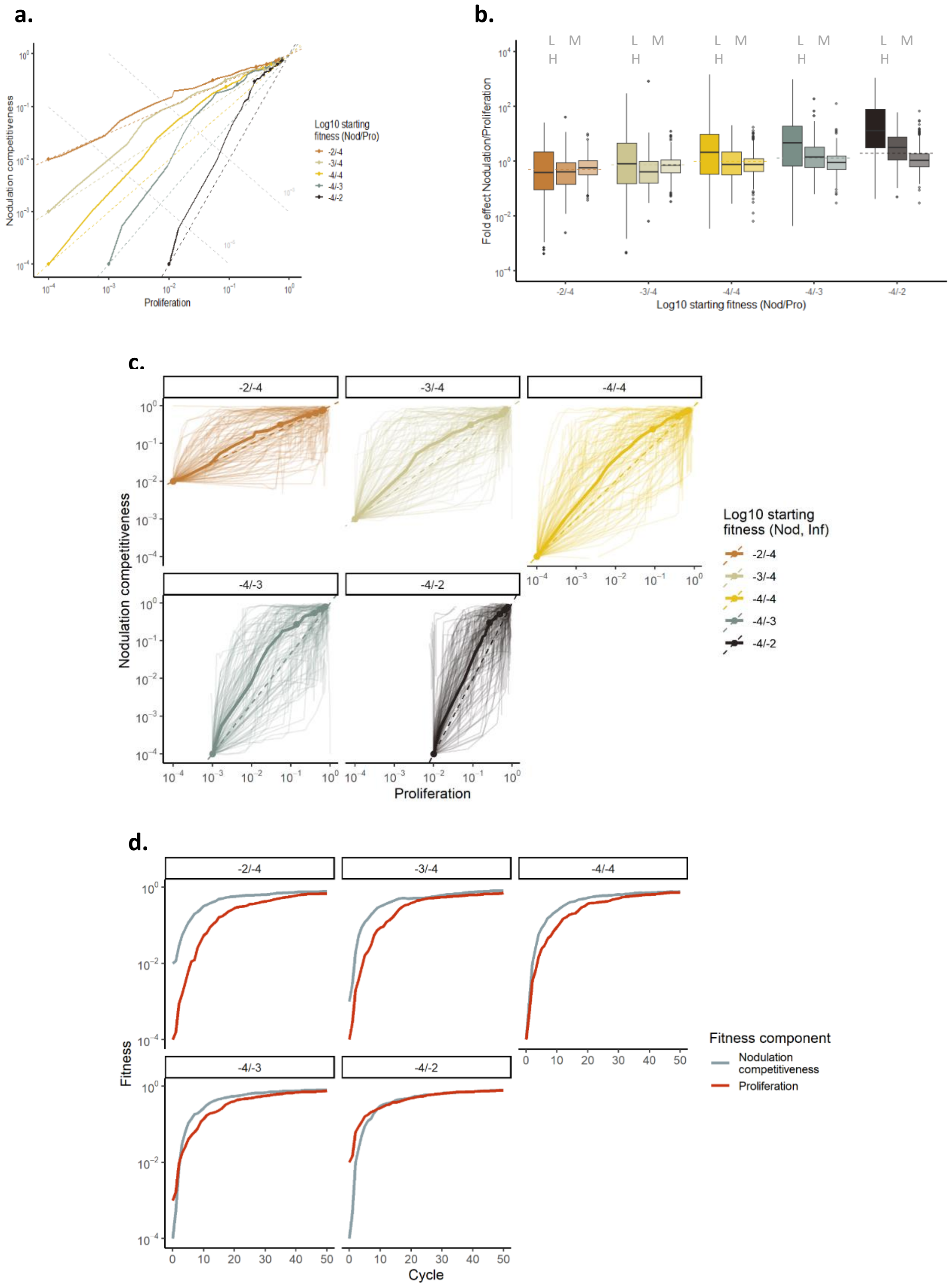


**Figure 6 – figure supplement 8:** Effect of the fitness of the ancestor on the relative strength of selection for nodulation competitiveness and proliferation, with a higher probability of beneficial mutations. a, b, as in Figure 4c,d (main text). c,d, as in Supplementary Figure 4a,b, with fitness trajectories shown for various combinations of nodulation competitiveness/proliferations fitness values in the ancestor ($10^{-2}/10^{-4}$), ($10^{-2}/10^{-4}$), ($10^{-4}/10^{-4}$), ($10^{-4}/10^{-3}$), and ($10^{-4}/10^{-2}$). Evolutionary parameters used were: k = l = 3 and m = 0.

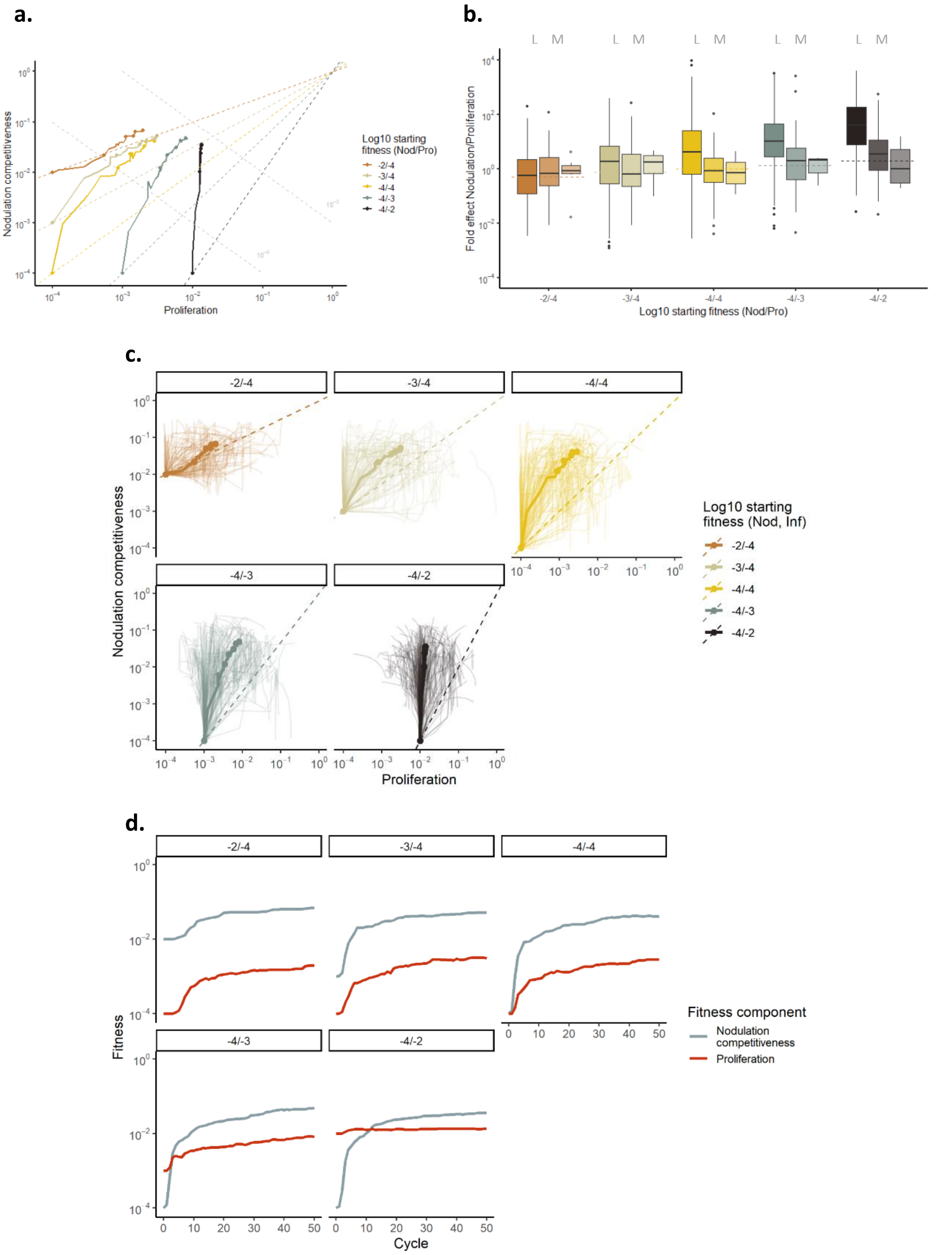


**Figure 6 – figure supplement 9:** Effect of the fitness of the ancestor on the relative strength of selection for nodulation competitiveness and proliferation, with a lower probability of beneficial mutations**.** a, b, as in Figure 4c,d (main text). c,d, as in Supplementary Figure 4a,b, with fitness trajectories shown for various combinations of nodulation competitiveness/proliferations fitness values in the ancestor ($10^{-2}/10^{-4}$), ($10^{-2}/10^{-4}$), ($10^{-4}/10^{-4}$), ($10^{-4}/10^{-3}$), and ($10^{-4}/10^{-2}$). Evolutionary parameters used were: k = l = 20 and m = 0.

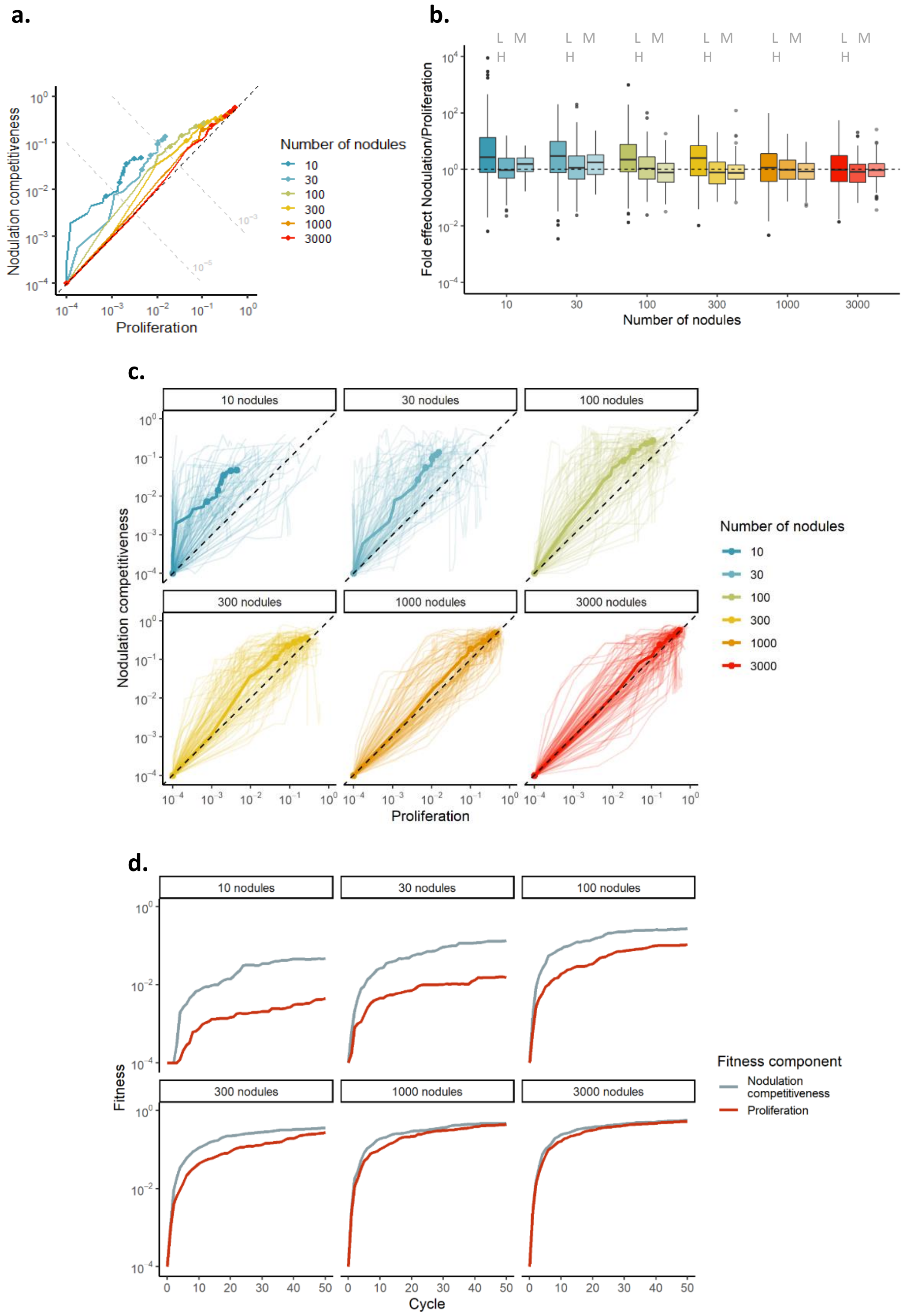


**Figure 6 – figure supplement 10:** Effect of the nodulation bottleneck on the relative strength of selection for nodulation competitiveness and proliferation under weak partial pleiotropy. a, b, as in Figure 4a,b (main text). c,d, as in Supplementary Figure 4a,b. Evolutionary parameters used were: k = l = 6 and m = 4.

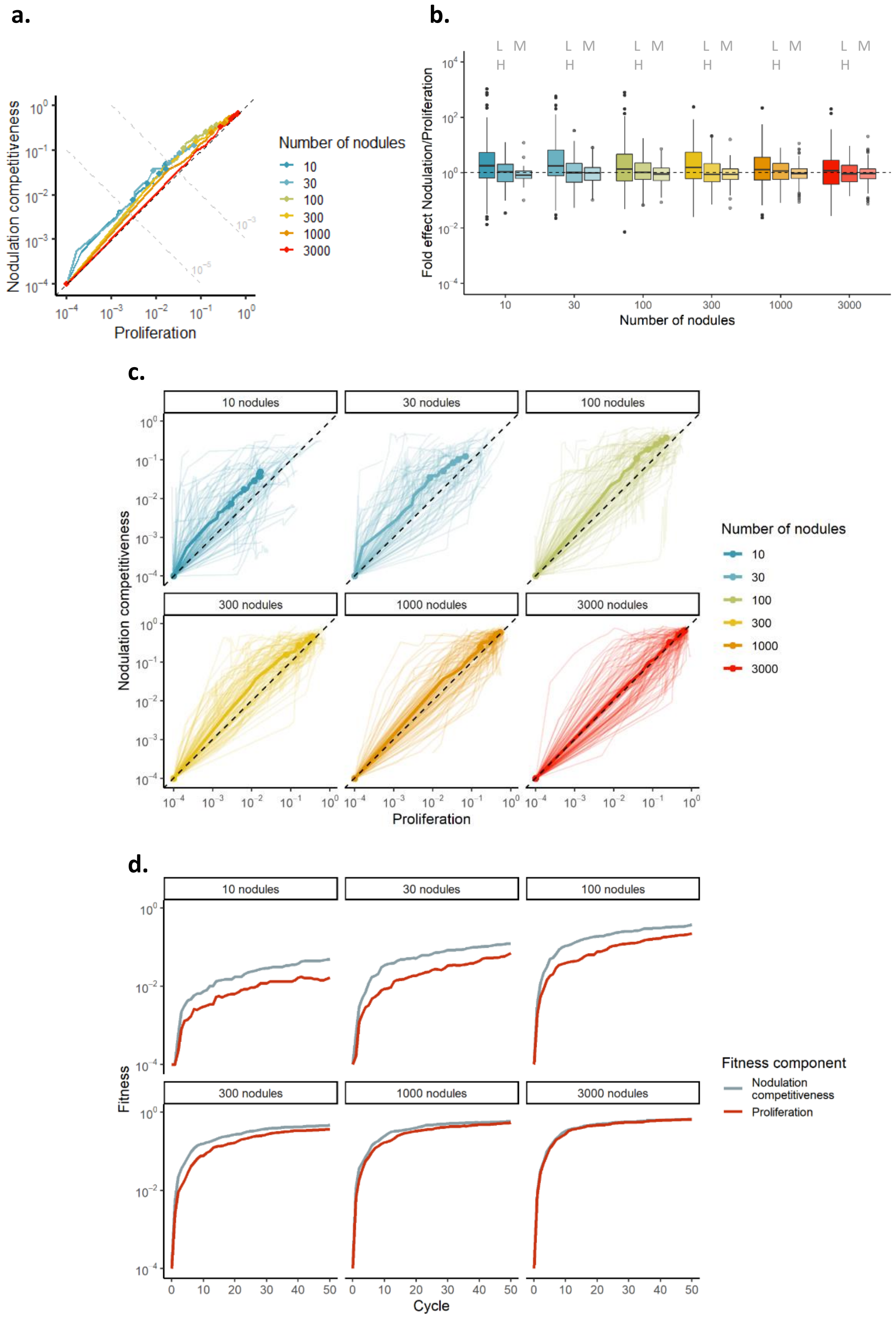


**Figure 6 – figure supplement 11:** Effect of the nodulation bottleneck on the relative strength of selection for nodulation competitiveness and proliferation under strong partial pleiotropy. a, b, as in Figure 4a,b (main text). c,d, as in Supplementary Figure 4a,b. Evolutionary parameters used were: k = l = 2 and m = 8.

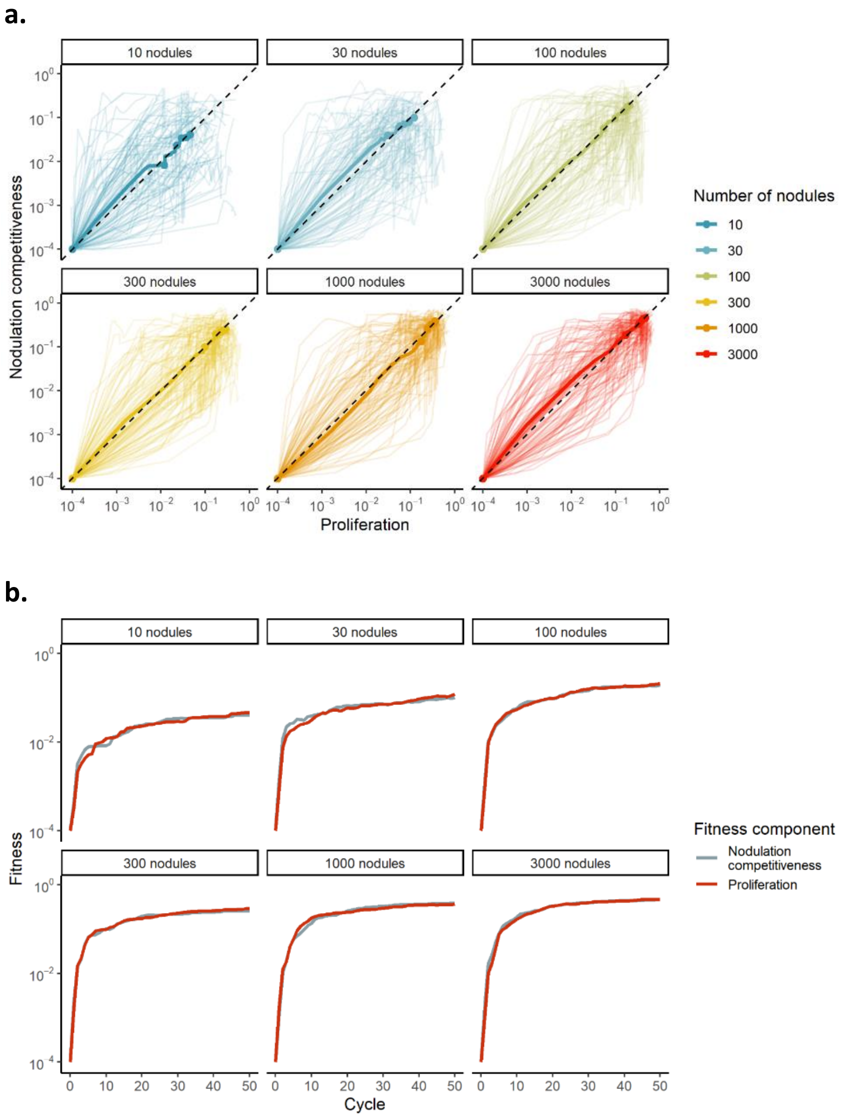


**Figure 6 – figure supplement 12:** Effect of the chronology of symbiotic events and the size of nodulation bottleneck on the relative strength of selection for nodulation competitiveness and proliferation. Same data as in Figure 4e,f (main text). a, b, as in Supplementary Figure 4a,b. Evolutionary parameters used were: k = l = 10 and m = 0.
